# Contrasting gene gain and loss shape metabolic diversification in the *Ralstonia solanacearum* species complex

**DOI:** 10.64898/2026.10.09.757798

**Authors:** Yingting Luo, Siqi Yang, Qingmei Liu, Xiaofan Zhou

**Author notes:** **Correspondence to**: Xiaofan Zhou. These authors contributed equally to this work (Yingting Luo, Siqi Yang).

## Abstract

Metabolic repertoires are continually remodeled by gene acquisition and loss, but how these contrasting processes jointly generate phenotypic diversity within bacterial species complexes remains poorly understood. Here, we elucidated the genetic and evolutionary basis of the classical biovar phenotypes of the *Ralstonia solanacearum* species complex (RSSC), which are defined by differential utilization of three hexitols (sorbitol, mannitol, and dulcitol) and three disaccharides (lactose, maltose, and cellobiose). Although this classification has been used for over half a century, the genetic determinants and evolutionary origins of these phenotypes have not been systematically resolved. Machine-learning analysis of genomic data from 250 RSSC strains with documented biovar phenotypes, combined with genome-wide association analysis and gene deletion, identified the *smoEFGK* gene cluster and flavin adenine dinucleotide (FAD)-dependent glucose dehydrogenase (FADGDH) as the major determinants of hexitol and disaccharide utilization, respectively. Comparative phylogenomics revealed fundamentally different evolutionary histories of these loci. An ancestral *smoEFGK* system was lost early in RSSC evolution and subsequently reacquired by the ancestor of phylotype I through horizontal transfer from a *Burkholderia*-related lineage, whereas vertically inherited FADGDH underwent recurrent, independent losses across multiple RSSC lineages. Extending the analysis to 1,082 RSSC strains further showed that disaccharide-utilization capacity was significantly depleted among monocot-associated strains after accounting for phylogenetic relatedness and geographic origin. Together, these findings reveal how contrasting trajectories of gene gain and loss can reshape bacterial metabolic repertoires, generate lineage-associated phenotypic diversity, and potentially contribute to ecological differentiation.

## Introduction

Metabolic capacity is a major dimension of bacterial phenotypic diversity and can change substantially over evolutionary time as gene repertoires expand and contract ^1^. New metabolic functions can arise through gene duplication followed by functional divergence ^2^, or through the acquisition of genes from other lineages by horizontal gene transfer ^3^, whereas gene loss can eliminate functions that have become dispensable or costly under particular ecological conditions ^4^. These processes can reshape metabolic repertoires in different ways and thereby contribute to phenotypic diversification. However, how gene gain and loss jointly generate metabolic diversity within bacterial species complexes remains less well understood. Addressing this question requires systems in which well-defined metabolic phenotypes can be linked to their genetic determinants and traced across an evolutionary radiation.

The *Ralstonia solanacearum* species complex (RSSC) provides an informative system for investigating these processes. RSSC comprises a phylogenetically diverse group of bacterial plant pathogens that causes bacterial wilt disease worldwide and collectively infect an exceptionally broad range of hosts spanning both monocot and eudicot lineages ^5,6,7^. RSSC is widely distributed across tropical and subtropical regions and also occurs in temperate environments ^7,8^, and can persist outside plant hosts in soil and aquatic environments for extended periods ^9–11^. Together, this broad host range, wide geographic distribution, and ability to occupy diverse environmental niches highlight the remarkable ecological versatility of the species complex. RSSC also exhibits substantial genomic plasticity, with mosaic genome architecture, extensive gene-content variation, and frequent horizontal gene acquisition documented across strains ^12–14^. These features make RSSC a particularly useful system for examining how genome variation translates into phenotypic diversity.

Metabolic capacity is central to the ecological breadth of RSSC. During its life cycle, RSSC transitions among environments that differ substantially in nutrient availability, including soil, the rhizosphere, and host xylem ^15,16,17^. Nutrient acquisition supports bacterial proliferation, while resources must also be allocated to processes required for host colonization and virulence ^18^. Accordingly, metabolic regulation and flexibility contribute to environmental persistence and successful colonization of plant hosts ^16,17^. One long-recognized manifestation of metabolic diversity within RSSC is the classical biovar system, established in the 1960s on the basis of differential capacity to oxidize three hexitols (mannitol, sorbitol, and dulcitol [galactitol]) and three disaccharides (lactose, maltose, and cellobiose) (Table 1) ^19,20^. The four classical biovars therefore represent the four possible combinations of two metabolic traits (utilization of hexitols and disaccharides). Biovar 1 strains utilize neither group of substrates, biovar 2 strains utilize disaccharides but not hexitols, biovar 3 strains utilize both substrate groups, and biovar 4 strains utilize hexitols but not disaccharides (Table 1). Subsequent phylogenetic analyses divided RSSC into four phylotypes, now classified into three species: *R. pseudosolanacearum* (phylotypes I and III), *R. solanacearum* (phylotype II), and *R. syzygii* (phylotype IV) ^21,22^. Although genome-based phylogenetic classification has superseded biovars for taxonomic purposes, the classical biovar system remains a simple and extensively documented readout of metabolic variation across RSSC lineages.

**Table 1.** Differentiation of RSSC biovars based on the ability to utilize three hexitols and three disaccharides.

| Substrates |  | Biovars |  |  |  |
| --- | --- | --- | --- | --- | --- |
|  |  | 1 | 2 | 3 | 4 |
| Hexitols | Sorbitol | - | - | + | + |
|  | Mannitol | - | - | + | + |
|  | Dulcitol | - | - | + | + |
| Disaccharides | Lactose | - | + | + | - |
|  | Maltose | - | + | + | - |
|  | Cellobiose | - | + | + | - |

-, no substrate utilization; +, substrate utilization

Despite more than half a century of biovar phenotyping, the genetic determinants and evolutionary origins of these metabolic differences have not been systematically resolved. An early comparative genomic study identified a 22-kb genomic region associated with the hexitol-positive phenotype ^23^. However, the specific genetic components responsible for the phenotype had not been functionally resolved, and the genetic basis of disaccharide utilization remained unknown. More broadly, it was unclear whether the two metabolic traits defining the classical biovars arose through similar or distinct evolutionary processes. Resolving their molecular basis therefore would provide a foundation for tracing the evolutionary histories of the underlying metabolic loci and for understanding how changes in gene repertoire generate phenotypic diversity among closely related bacterial lineages.

In this study, we combined genomic and biovar data from 250 RSSC strains with machine-learning, genome-wide association, and experimental approaches to identify the major loci associated with the two metabolic traits (Fig. 1A). We then reconstructed the evolutionary histories of these loci across RSSC and related bacteria, and extended the analysis to a broader collection of 1,082 RSSC strains to examine whether variation in metabolic capacity was associated with host affiliation. This integrated framework allowed us to connect the molecular basis of classical biovar phenotypes with broader patterns of genome evolution and ecological differentiation across the species complex.

**Figure 1.**
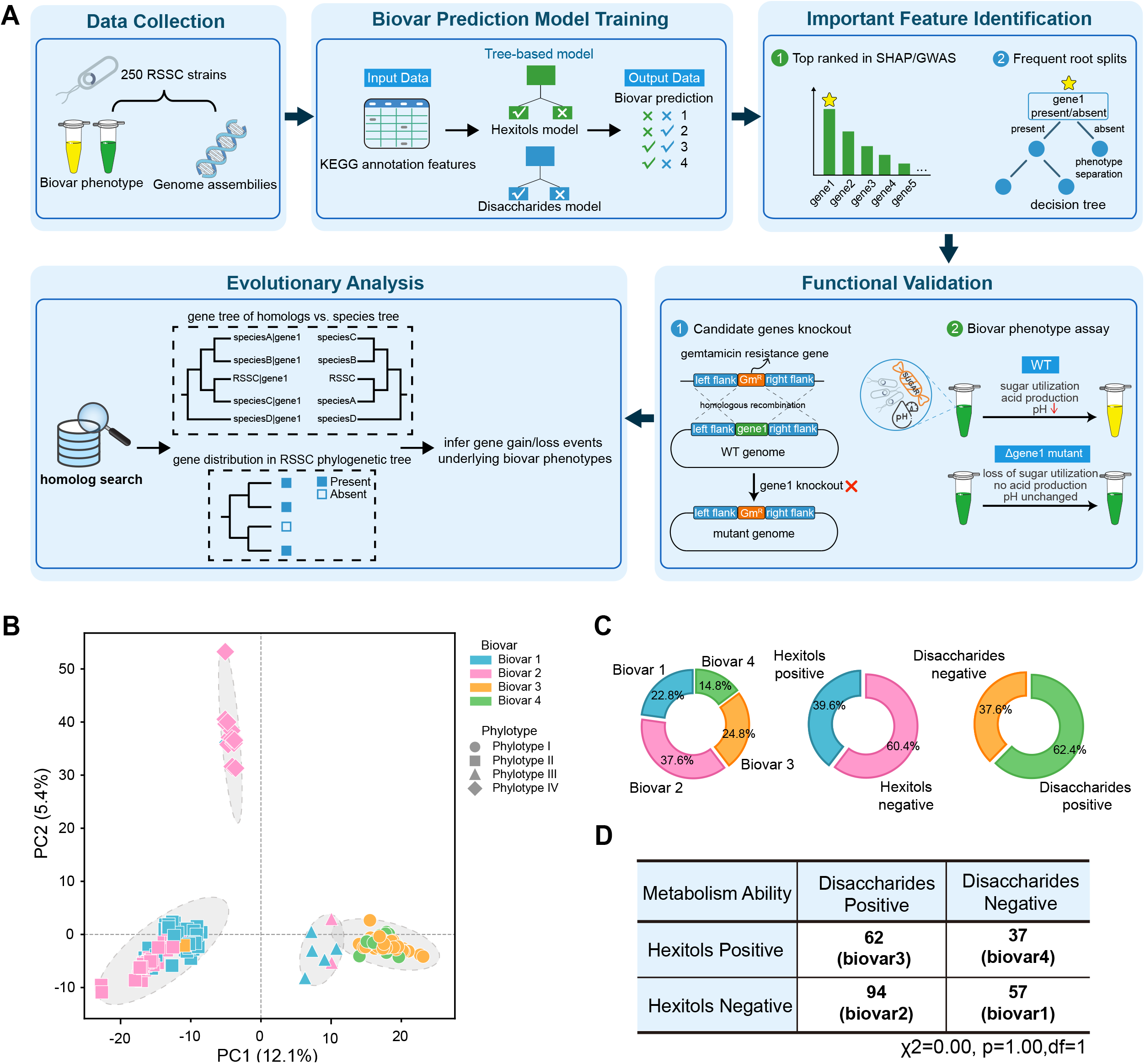
Overview of the study design and the 250-strain RSSC dataset. (A) Overview of the study framework. (B) PCA of KEGG Orthology (KO) copy-number profiles of the 250 RSSC strains. Points are colored by biovar and shaped by phylotype. (C) Distribution of biovar classes and the two metabolic traits in the 250-strain dataset. The three doughnut charts show the proportions of biovars, hexitol utilization phenotypes, and disaccharide utilization phenotypes, respectively. (D) Contingency table for chi-square analysis of the association between hexitol and disaccharide utilization phenotypes.

## Results

### A 250-strain dataset spanning all RSSC phylotypes and classical biovars

We assembled a dataset of 250 RSSC strains with documented biovar phenotypes and available genome assemblies (Table S1). Genome-wide profiles of KEGG Orthology (KO) counts derived from protein annotations were used to characterize variation in functional gene repertoire across strains. Principal component analysis (PCA) of these functional profiles revealed clear clustering corresponding to the four phylotypes, indicating substantial phylogenetic structure in genomic composition (Fig. 1B). Biovar 3 and 4 strains were also separated from biovar 1 and 2 strains, consistent with the known restriction of biovar 3 and biovar 4 to phylotype I, whereas biovar 1 and biovar 2 occur in phylotypes II–IV ^21^.

The four biovar classes were well represented in the dataset, with biovar 2 accounting for the largest proportion (94 strains, 37.6%), followed by biovar 3 (62 strains, 24.8%), biovar 1 (57 strains, 22.8%) and biovar 4 (37 strains, 14.8%) (Fig. 1C and 1D). In terms of metabolic traits, hexitol-positive and disaccharide-positive strains comprised 39.6% and 62.4% of the dataset, respectively (Fig. 1C), such that both positive and negative states of each trait were represented by substantial numbers of strains, providing a reasonable basis for downstream genotype–phenotype analyses. A chi-square analysis detected no significant association between the two metabolic traits (P = 1.00; Fig. 1D), raising the possibility that they are governed by distinct genetic determinants.

### Genome-wide functional profiles accurately predict the metabolic traits underlying biovar classification

Because hexitol and disaccharide utilization represent two distinct components of the biovar classification, we modeled the two metabolic traits separately and subsequently integrated their predictions to infer the biovar of each strain (Fig. 2A). Gene counts for each KO identifier in each of the 250 RSSC strains were used as input features, encompassing 2,566 distinct KOs (Table S2). For each metabolic trait, we compared random forest and gradient boosting tree classifiers and selected the better performing classifier based on overall cross-validation performance (Fig. S1A). The selected hexitol-utilization model used a random forest classifier, whereas the disaccharide-utilization model used a gradient boosting tree classifier.

**Figure 2.**
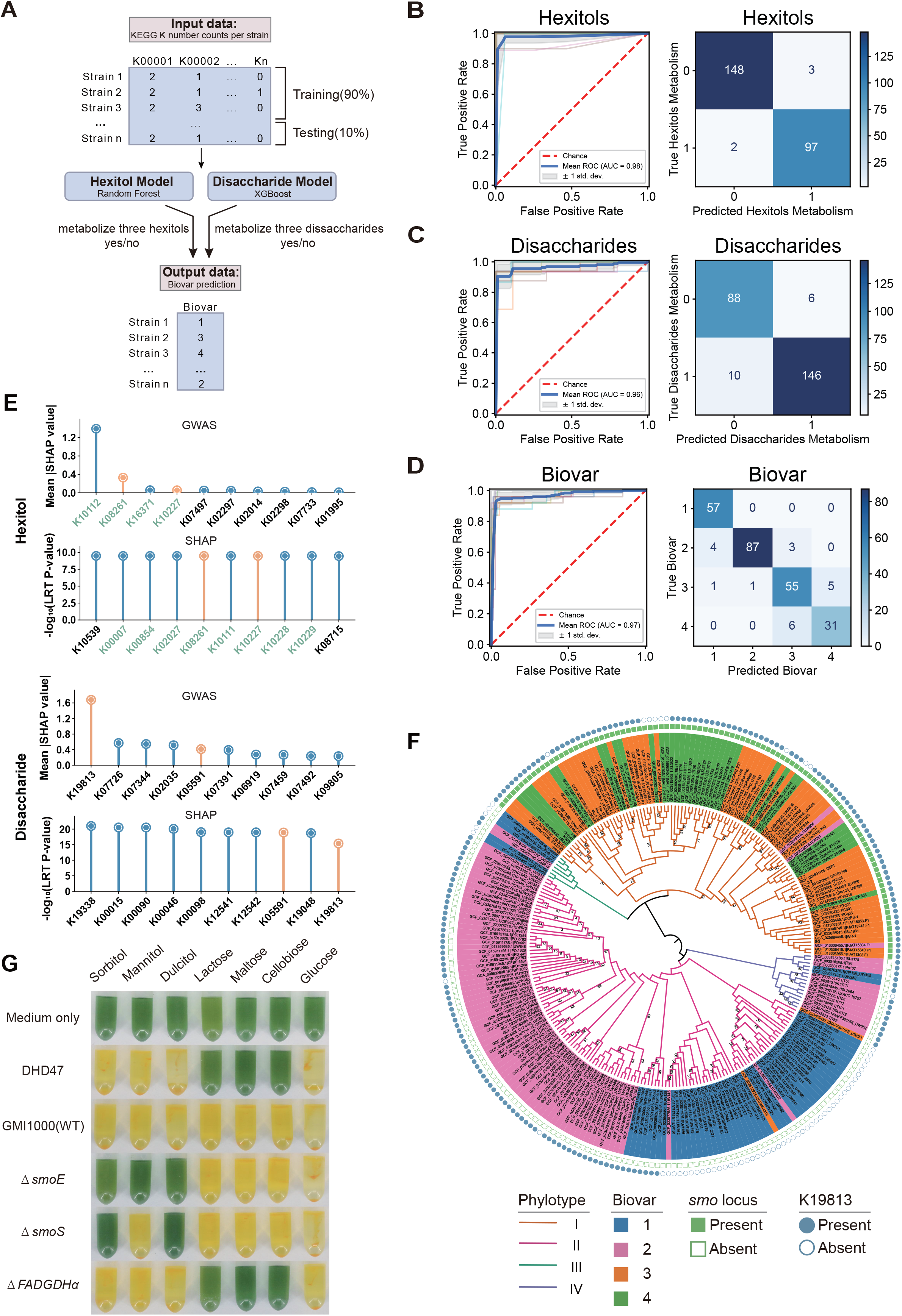
Prediction and genetic dissection of the metabolic traits underlying RSSC biovars. (A) Workflow for the models predicting the two metabolic traits underlying biovar classification. Genomewide KO copy-number profiles of the 250 RSSC strains were used as input features. Model performance was evaluated using 10-fold cross-validation, in which the model was trained on nine folds and tested on the one remaining fold in each of the 10 iterations. Separate models were trained to predict hexitol and disaccharide utilization, and their predictions were subsequently integrated to infer the biovar of each strain. (B-D) Receiver operating characteristic (ROC) curves and confusion matrices for the hexitol-utilization model (B), disaccharide-utilization model (C), and integrated biovar model (D). ROC curves show the relationship between the true positive rate and false positive rate across different classification thresholds. The confusion matrices display the distribution of correct and incorrect predictions, with diagonal cells representing correctly classified strains. Both the ROC curves and the confusion matrices were derived from 10-fold cross-validation. (E) Genomic features prioritized by model interpretation and genome-wide association analysis. The top 10 features identified by SHAP analysis of the machine-learning models and the top 10 phenotype-associated features identified by genome-wide association analysis are shown separately for hexitol and disaccharide utilization. Features identified by both approaches are marked with orange lollipop symbols. Features associated with the *smo* locus are highlighted in green font. (F) Phylogenetic tree of 250 strains with annotations for phylotype, biovar, and the presence or absence of the *smo* locus and K19813, which corresponds to the FADGDH α-subunit. Bootstrap values below 95 are labeled on the branches. The tree is rooted at the midpoint. (G) Biovar assays of the wild type strain and gene deletion mutants. Uninoculated medium was included as a blank control. *R. pseudosolanacearum* DHD47 (biovar 4) was used as a negative control for disaccharide utilization, and *R. pseudosolanacearum* GMI1000 was used as the wild type control. Δ*smoE*, Δ*smoS*, and *ΔFADGDHa* represent gene deletion mutants constructed in the GMI1000 background. A positive reaction is indicated by a color change of the bromothymol-blue indicator from green to yellow due to medium acidification.

Both selected models showed high predictive performance. The hexitol-utilization model achieved an accuracy of 98.0% and a balanced accuracy of 98.1%, while the disaccharide-utilization model achieved an accuracy of 93.7% and a balanced accuracy of 93.8% (Fig. S1A). Integrating predictions from the two models for biovar classification achieved an overall accuracy of 92.1% and a balanced accuracy of 92.4% (Fig. S1A). The mean areas under the receiver operating characteristic (ROC) curves (AUCs) were 0.98, 0.96 and 0.97 for hexitol utilization, disaccharide utilization, and integrated biovar classification, respectively (Fig. 2B–D). Confusion matrices also showed few misclassifications across all three tasks (Fig. 2B–D). Precision–recall curves and additional evaluation metrics, including F1 score and Matthews correlation coefficient (MCC), showed similar strong performance (Fig. S1). Together, these results indicate that genome-wide functional profiles contain substantial predictive information for the metabolic traits underlying RSSC biovar classification.

### *smo* locus and FADGDH are major determinants of hexitol and disaccharide utilization

To identify genomic features contributing to the two metabolic traits, we used SHapley Additive exPlanations (SHAP) analysis to quantify the contributions of individual KO features to model predictions. For hexitol utilization, the four highest-ranked features, K10112, K08261, K16371, and K10227, corresponded to genes within the same genomic locus (Fig. 2E and Fig. 3A). Genome-wide association (GWAS) analysis also identified KO features corresponding to genes from this locus as being associated with hexitol utilization, providing independent support for their relationship with the phenotype (Fig. 2E). This genomic locus corresponds to the *smo* (sorbitol mannitol oxidation) locus, previously characterized in *Sinorhizobium meliloti* ^24,25^. The *smoEFGK* genes encode components of an ABC transporter that mediate the uptake of sorbitol, mannitol and dulcitol, while neighboring genes encode enzymes (e.g., dehydrogenases) that catalyze their subsequent metabolism (Fig. 3A). Notably, this genomic locus is found only in phylotype I in the 250-strain dataset, closely matching the distribution of the hexitol utilization phenotype (Fig. 2F).

**Figure 3.**
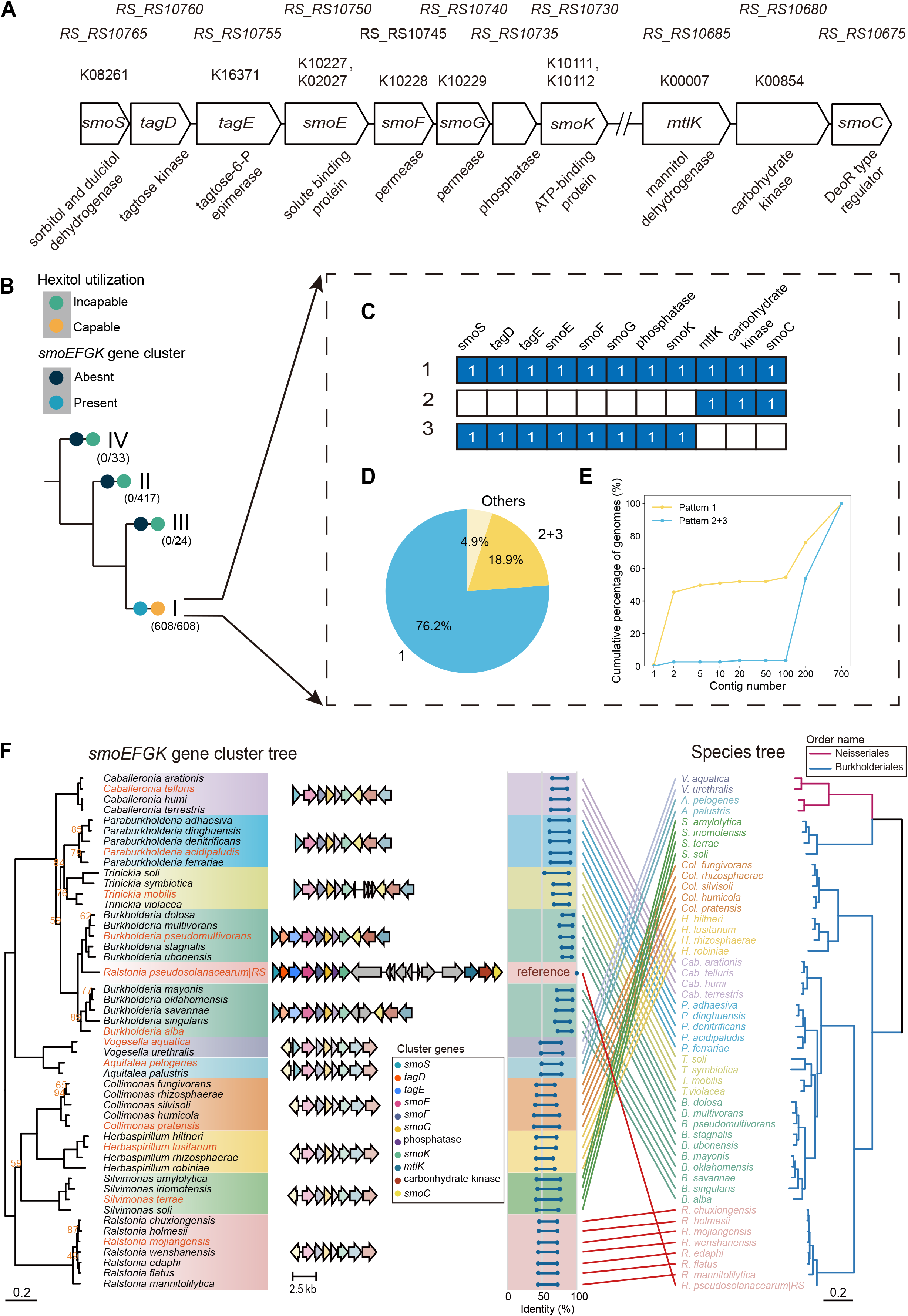
Distribution and horizontal acquisition of the *smo* locus in RSSC. (A) Genomic organization of the *smo* locus in *R. pseudosolanacearum* GMI1000. From top to bottom, the four rows indicate GMI1000 gene locus tags, KEGG Orthology identifiers, gene symbols, and predicted gene functions, respectively. The *smo* locus is primarily composed of an ABC transporter together with associated dehydrogenases. The ABC transporter components encoded by *smoE*, *smoF*, *smoG*, and *smoK* mediate the uptake of sorbitol, mannitol and dulcitol. Following transport, the dehydrogenases SmoS and MtlK catalyze substrate oxidation. SmoS oxidizes sorbitol to fructose and dulcitol to tagatose, after which tagatose is phosphorylated by TagD and subsequently epimerized by TagE to produce fructose-6-phosphate 25. MtlK catalyzes the oxidation of mannitol to fructose. (B) Distribution of the *smo* locus across RSSC phylotypes I-IV. Numbers in parentheses indicate the number of genomes carrying the *smo* locus relative to the total number of genomes analyzed in each phylotype. (C) Major patterns of the *smo* locus among phylotype I assemblies. Pattern 1 contains the locus as a continuous genomic region, whereas patterns 2 and 3 represent complementary portions of the locus recovered separately in assemblies. (D) Frequencies of *smo* locus patterns in phylotype I genomes. Pattern 2 + 3 denotes assemblies in which both patterns 2 and 3 were recovered. (E) Cumulative distributions of contig numbers for genomes assigned to pattern 1 and pattern 2+3. The x-axis is shown on a logarithmic scale. (F) Comparison of the *smo* locus phylogeny with the corresponding species phylogeny. The locus tree was inferred from a concatenated alignment of seven conserved genes (*smoS*, *smoE*, *smoF*, *smoG*, *smoK*, *mtlK*, and *smoC)* from RSSC and related bacteria. Background colors represent different bacterial families. For each family, a representative gene cluster was visualized using clinker, and the corresponding species names in the tree are colored orange. Gene transparency reflects the sequence identity to the corresponding genes of *R. pseudosolanacearum* strain RS. The blue lollipop plot indicates the range of sequence identities (lowest to highest) across genes within each cluster. The species in the tree belong to two orders, Neisseriales and Burkholderiales. Bootstrap values below 95 are labeled on the branches. All bootstrap values in the species tree are at least 95.

For disaccharide utilization, SHAP and GWAS analyses converged on two features, K19813 and K05591 (Fig. 2E). K19813 corresponds to the α-subunit of flavin adenine dinucleotide (FAD)-dependent glucose dehydrogenase (FADGDH), while K05591 corresponds to the ATP-dependent RNA helicase DbpA. K19813 showed a substantially higher mean absolute SHAP value than other candidate features and was selected as the root split in 12 of the 100 trees comprising the disaccharide model, indicating a strong contribution to model predictions. In the 250-strain RSSC dataset, the absence of K19813 annotation was strongly associated with the inability to utilize the three disaccharides (Fig. 2F). Specifically, among the 55 strains without K19813 annotation, 52 were unable to utilize lactose, maltose, or cellobiose. The remaining three strains (UW145, UW272 and CRMRs218) all had biovar assignments that were phylogenetically incongruent with their respective sister clades (Fig. 2F), raising the possibility of phenotype annotation discrepancies. Conversely, the majority of strains carrying K19813 were able to utilize the three disaccharides (153/195). Taken together, these observations suggest a general requirement for FADGDH in disaccharide utilization across RSSC.

We next tested these candidate genes experimentally in *R. pseudosolanacearum* GMI1000 (Fig. 2G). Under the biovar assay conditions, deletion of *smoE* abolished the utilization of all three hexitols; deletion of *smoS* abolished sorbitol and dulcitol utilization but retained mannitol utilization; and deletion of the gene encoding FADGDH *α*-subunit (*FADGDHα*) abolished the utilization of all three disaccharides, demonstrating that FADGDH is required for disaccharide utilization in GMI1000. Altogether, these deletion phenotypes provide functional support for the roles of the SmoEFGK system and FADGDH in the corresponding metabolic traits.

### The *smo* locus was horizontally acquired in the ancestor of RSSC phylotype I

The typical *smo* locus in phylotype I, as represented by *R. pseudosolanacearum* GMI1000, comprises genes involved in hexitol transport and metabolism, including the *smoEFGK* ABC transporter genes, the dehydrogenase genes *smoS* and *mtlK*, and several associated metabolic genes, including *tagD*, *tagE*, a phosphatase gene, a carbohydrate kinase gene, and *smoC* (Fig. 3A). Most of these genes form a compact genomic region, with the *mtlK*–*smoC* portion separated from the upstream block by eight intervening genes. We next examined the distribution and conservation of the *smo* locus across publicly available RSSC genome assemblies. The *smo* locus was detected in all 608 analyzed phylotype I genomes, whereas it was not detected in any strains belonging to phylotypes II–IV (Fig. 3B).

Based on how *smo*-locus genes were recovered in phylotype I genome assemblies, three main patterns were identified (Fig. 3C and 3D). Pattern 1 contained the complete *smo* locus as a single continuous region, whereas patterns 2 and 3 represented two compact gene blocks that always co-occurred in the same assembly. Because these two blocks correspond to complementary portions of the complete locus in GMI1000, separated by eight intervening genes (Fig. 3A and 3C), we examined whether their separation reflected a genuine genomic arrangement or fragmented genome assembly. Because RSSC genomes comprise two replicons ^13^, (near-)complete assemblies are therefore expected to contain approximately two contigs. Assemblies with two contigs accounted for 44.5% of pattern 1 strains but only 2.6% of pattern 2 + 3 strains. Moreover, 96.5% of pattern 2 + 3 strains (111/115) were assembled into more than 100 contigs and all of these assemblies had at least one of the two gene blocks located near a contig boundary (Fig. 3E; Table S3). These results suggest that the separation observed in patterns 2 and 3 is primarily attributable to assembly fragmentation rather than distinct biological organization. Together, pattern 1 and pattern 2 + 3 accounted for 95.1% of phylotype I assemblies, indicating strong conservation of the locus within this lineage (Fig. 3D).

We next investigated the evolutionary origin of the *smo* locus in phylotype I. Phylogenetic analysis of individual locus genes consistently placed homologs from RSSC phylotype I (represented by *R. pseudosolanacearum* strain RS) nested within clades of *Burkholderia* sequences, rather than with homologs from other *Ralstonia* species (Fig. S2). Approximately unbiased (AU) topology tests rejected constrained topologies enforcing monophyly of *Ralstonia* homologs for most analyzed genes, including *smoE*, *smoF*, *smoK*, *mtlK*, and *smoC* (AU test, P < 0.05; Table S4). A phylogeny reconstructed from the concatenated alignment of seven *smo*-locus genes recovered the same relationship, in which the locus from phylotype I grouped with homologous loci from *Burkholderia* species, whereas loci from other *Ralstonia* species formed a separate lineage (Fig. 3F). Moreover, the *smo* locus of phylotype I showed higher sequence similarity and more similar gene organization to homologous loci of *Burkholderia* than to those of other *Ralstonia* species, including the linkage of *tagD* and *tagE* to the core *smo*-locus genes (Fig. 3F). Together, these results support horizontal acquisition of the *smo* locus in the ancestor of RSSC phylotype I from a *Burkholderia*-related lineage.

Homologous *smo* loci are present in non-RSSC *Ralstonia* species but absent from RSSC phylotypes II–IV (Fig. 3F). This broader phylogenetic distribution is consistent with a scenario in which an ancestral *smo*-like system was lost early in RSSC evolution, followed by re-acquisition of a homologous locus in the ancestor of phylotype I through horizontal gene transfer (HGT).

### FADGDH underwent recurrent losses across RSSC lineages

Deletion of the gene encoding FADGDH α-subunit abolished the utilization of all three disaccharides, but whether RSSC FADGDH (RsGDH) could directly recognize these substrates remained unclear. FADGDH from *Burkholderia cepacia* (BcGDH) has been reported to exhibit broad substrate specificity, including activity toward maltose ^26^. FADGDH consists of three subunits: a catalytic α-subunit containing an FAD cofactor and a 3Fe-4S iron-sulfur cluster, a β-subunit that mediates electron transfer through three heme c groups, and a γ-subunit functioning as a hitchhiker protein that facilitates the transport of folded proteins from the cytoplasm to the periplasm via the twin-arginine transport (TAT) pathway ^27^. The α-subunit of FADGDH contains the substrate-binding pocket and constitutes the primary catalytic component of the enzyme. In GMI1000, this subunit shares 81% protein sequence identity with BcGDH, and the characterized active-site residues are fully conserved (Fig. S3). Homology modeling of RsGDH using BcGDH as the template produced a highly similar structure (root-mean-square deviation [RMSD] = 0.098 Å) (Fig. S4A).

Molecular docking of D-glucose to RsGDH revealed a substrate-binding orientation broadly consistent with that observed for D-glucono-1,5-lactone (LGC) in the BcGDH crystal structure ^27^ (Fig. 4A). Five of the seven hydrogen-bonding glucose–residue pairs observed in BcGDH (O1–His476, O1–Asn519, O2–Asn475, O2–Asn474, O3–Asn474, O6–His363, and O6–Glu341) were reproduced in the unconstrained RsGDH docking model (Fig. 4A; Fig. S4B). Moreover, the substrate-binding cavity of RsGDH is open toward the protein surface, providing sufficient space to accommodate lactose, maltose, and cellobiose when their reducing-end glucose residues were positioned in the catalytic pocket (Fig. 4B). These structural analyses provide a plausible basis for the ability of RsGDH to accommodate all three disaccharides.

**Figure 4.**
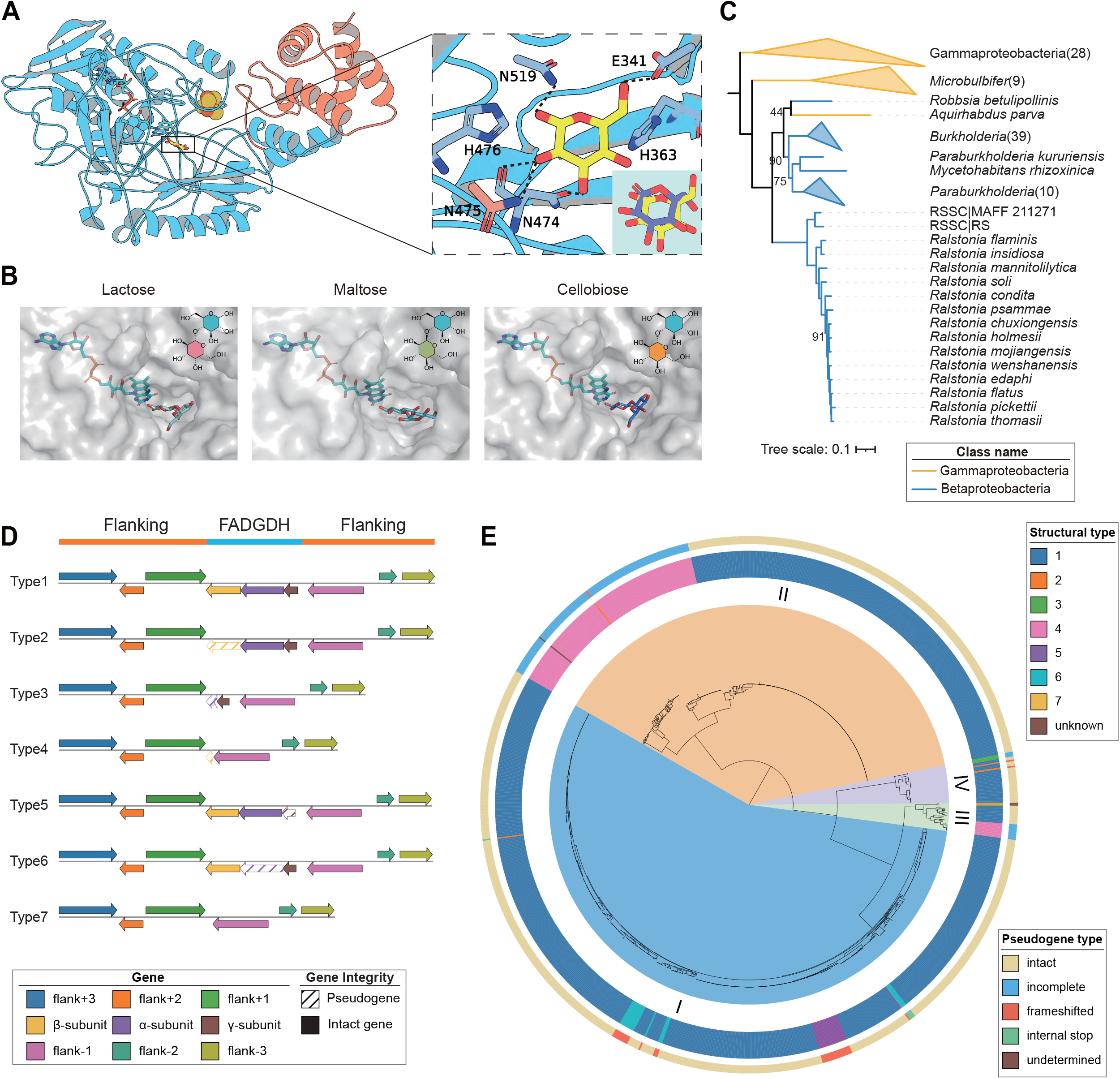
Structural modeling and evolutionary history of FADGDH in RSSC. (A) Homology-based structural model of the RSSC FADGDH αγ complex with docked glucose. The structure was constructed by homology modeling using *Burkholderia cepacia* FADGDH (PDB ID: 8GRJ) as the template. The α-subunit and γ-subunit are colored blue and orange, respectively. Glucose is shown as yellow sticks, the FAD cofactor as cyan sticks, and the iron-sulfur cluster (3Fe-4S) as yellow and orange spheres. The enlarged panel on the right shows the docked glucose and labels six active-site residues identified in the BcGDH crystal structure. Five of the seven corresponding hydrogen-bonding interactions were reproduced in the RsGDH docking model and are indicated by black dashed lines. The lower-right inset shows an overlay of the binding poses of docked glucose (yellow) and D-glucono-1,5-lactone (purple) from the *B. cepacia* FADGDH crystal structure. (B) Substrate-binding pocket views of RsGDH in complex with superimposed disaccharide substrates. The chemical structure of each disaccharide is shown above the corresponding panels, with the reducing-end glucosyl moiety highlighted in blue. Each disaccharide was positioned by pair fitting the C1-C6 atoms of its reducing end glucosyl moiety to those ofD-glucono-1,5-lactone in the *B. cepacia* FADGDH crystal structure. The resulting protein-ligand complexes were subsequently subjected to energy minimization. (C) Maximum-likelihood phylogeny of the FADGDH gene cluster across *Ralstonia* and closely related species, inferred from the concatenated protein sequences of the three FADGDH subunits. Bootstrap values below 95 are labeled on the branches. The tree was midpoint-rooted. Triangles denote collapsed clades, with the number of species shown in parentheses. (D) Representative organization types of the FADGDH gene cluster, with conserved flanking genes included. Type 1 represents the intact FADGDH gene cluster, while types 2-7 represent distinct forms of degeneration or loss. Pseudogenes are shown as diagonally hatched arrows. Representative genome accessions of the seven types are GCF_001373295.1 (type 1), GCF_003515165.1 (type 2), GCF_054491955.1 (type 3), GCF_001373255.1 (type 4), GCF_048285355.1 (type 5), GCF_015912175.1 (type 6), GCF_006088755.1 (type 7). (E) Phylogenetic tree of 1,082 RSSC strains annotated with FADGDH structural types. Phylotypes I-IV are labeled on the tree. Unknown structural types result from fragmented genome assemblies. “Incomplete” indicates a large sequence deletion within the locus.

We next reconstructed the evolutionary history of FADGDH using concatenated protein sequences of the three subunits from *Ralstonia* and related bacteria (Fig. 4C). Unlike the horizontally acquired *smo* locus, all *Ralstonia* FADGDH homologs formed a monophyletic lineage, consistent with vertical inheritance of FADGDH from a common ancestor of *Ralstonia*. Within the 250-strain dataset, FADGDH absence occurred in multiple phylogenetically separated lineages, suggesting recurrent loss during RSSC diversification (Fig. 2F). However, presence–absence patterns alone could not distinguish shared ancestral losses from independent loss events or reveal the mutational processes underlying FADGDH degeneration. We therefore expanded the analysis to 1,082 publicly available RSSC genome assemblies and examined structural remnants of the FADGDH locus.

Conserved genomic regions flanking the FADGDH gene cluster allowed us to identify FADGDH pseudogene remnants and classify cluster structures into seven distinct types (Fig. 4D and Fig. S5). These types differed in which of the α-, β-, and γ-subunit genes remained intact, had become pseudogenized, or were no longer detectable. Mapping these structural types onto the 1,082-strain RSSC phylogeny revealed FADGDH loss in all four phylotypes (Fig. 4E). In phylotype II, FADGDH losses were concentrated within two large monophyletic clades and were almost exclusively classified as type 4, suggesting two ancestral FADGDH losses in this phylotype. In contrast, the scattered distributions of distinct loss types in phylotypes I, III, and IV indicated multiple more recent loss events. Phylotypes III and IV each contained two different loss types, consistent with at least two independent loss events in each lineage. In phylotype I, types 5 and 6 each occurred in multiple separate parts of the phylogeny. Moreover, we found two distinct forms of type 6 associated with different disruptive mutations in the FADGDH α-subunit—a frameshift in one lineage and an internal stop codon in the other—providing strong evidence that similar patterns of degeneration arose independently (Fig. S6). Together, these results indicate that an ancestrally inherited FADGDH system has been recurrently lost during RSSC diversification.

### Disaccharide utilization is depleted among monocot-associated RSSC strains

To examine whether variation in the two metabolic traits was associated with ecological source or host affiliation, we analyzed a broader collection of 1,082 RSSC strains for which genome assemblies and phylogenetic relationships were available (Fig. 5A). Broad isolation-source information was available for 1,035 strains, including 852 plant-associated, 106 soil/rhizosphere-associated, and 77 water-associated strains. The 852 plant-associated strains could be further assigned to host clade, family, and species. Of the original 250 strains used for model training, 224 were represented in this expanded 1,082-strain dataset and therefore had experimentally documented hexitol- and disaccharide-utilization phenotypes; phenotypes for the remaining 858 strains were inferred using the corresponding machine-learning models. Detailed strain metadata and phenotype assignments are provided in Table S5.

**Figure 5.**
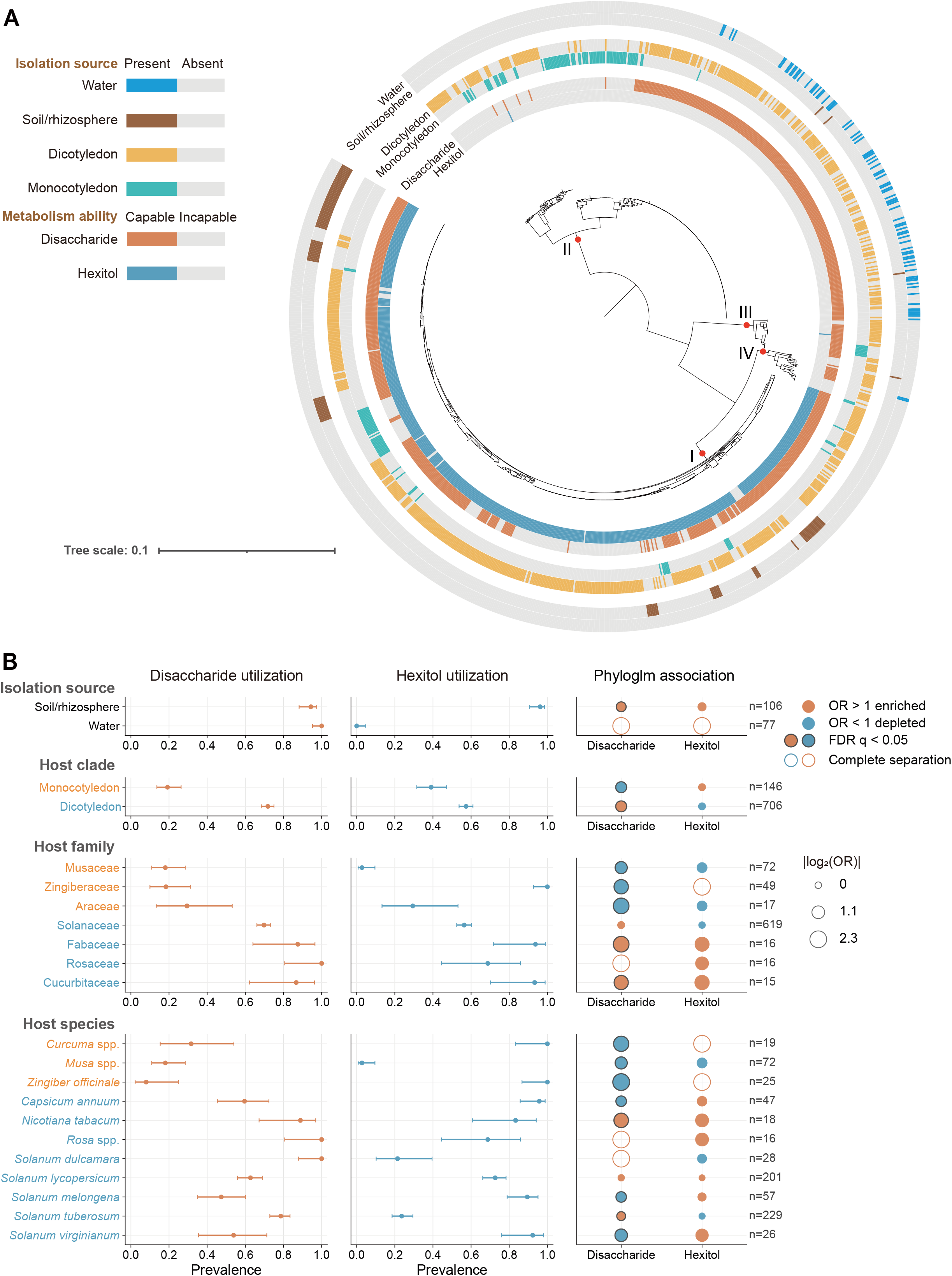
Ecological associations of hexitol- and disaccharide-utilization capacity across RSSC. (A) Phylogenetic distribution of isolation source, host clade, and metabolic capacity across 1,082 RSSC strains. Phylotypes are labeled on the branches. (B) Prevalence and phylogenetically adjusted associations of metabolic capacities with isolation source, host clade, host family and host species. Forest plots show observed prevalence estimates and Wilson 95% confidence intervals for each metabolic capacity. Bubble plots summarize odds ratios estimated using Ives-Garland phylogenetic logistic regression. Orange and blue bubbles indicate enrichment (OR > 1) and depletion (OR < 1), respectively, with bubble size proportional to |log_2_(OR)|. Black outlines indicate associations with an FDR q < 0.05. Open circles indicate complete separation, where the corresponding metabolic trait was either present in all strains or absent from all strains in the focal group; these cases were excluded from FDR correction and OR estimates were not interpreted. Sample sizes are shown on the right. Orange and blue labels indicate monocot and dicot hosts, respectively. Only focal categories with at least 10 strains were included in the association analyses and displayed in this plot. Hosts labeled “spp.” indicate that the NCBI metadata could not be resolved to a specific species.

We first tested associations between the two metabolic traits and ecological categories using Ives–Garland phylogenetic logistic regression, thereby accounting for shared evolutionary history among strains. The two traits showed markedly different association patterns. Hexitol utilization was strongly concentrated within phylotype I (Fig. 5A), and none of its apparent associations with isolation source or host categories remained significant after phylogenetic correction (all FDR q > 0.05; Fig. 5B). In contrast, disaccharide utilization retained significant associations with host affiliation. The clearest pattern occurred at the host-clade level: monocot-associated strains had a substantially lower prevalence of disaccharide utilization than dicot-associated strains (19.2% versus 71.8%), and this depletion remained significant after accounting for phylogenetic relatedness (phyloglm OR = 0.598, 95% CI = 0.402–0.887, FDR q = 0.01; Fig. 5B).

We next evaluated whether this monocot–dicot association was robust to alternative treatments of phylogenetic non-independence (Fig. S7). Logistic regression incorporating phylogenetic principal-coordinate/multidimensional-scaling (PC/MDS) axes recovered the same depletion of disaccharide utilization among monocot-associated strains (OR = 0.182, FDR q = 2.00 × 10⁻¹⁰). A phylogenetic generalized linear mixed model incorporating a tree-derived covariance matrix likewise recovered the same direction of effect (OR = 0.001). As a complementary evolutionary test, Pagel’s discrete correlated-evolution analysis significantly favored the dependent over the independent model for host clade and disaccharide utilization (ΔAIC = 95.41, bootstrap P = 0.002), supporting non-independent evolution of these two binary traits (Fig. S7).

Because metabolic phenotypes for most strains in the expanded dataset were inferred computationally, we repeated the phylogenetic logistic regression using only strains with experimentally supported disaccharide-utilization phenotypes. The depletion of disaccharide utilization among monocot-associated strains remained significant in this experimental-only analysis (phyloglm OR = 0.188, FDR q = 0.0013), showing that the association was also detectable independently of machine-learning-predicted phenotypes (Fig. S7). We further examined whether geographic structure contributed to the observed association by including geographic continent as an additional covariate in the phylogenetic logistic regression. The monocot-associated depletion remained significant after this adjustment (phyloglm OR = 0.621, FDR q = 0.0142; Fig. S7).

Consistent with the broad host-clade pattern, strains associated with the monocot families Musaceae, Zingiberaceae, and Araceae, as well as those associated with *Curcuma* spp., *Musa* spp., and *Zingiber officinale*, showed significantly reduced disaccharide-utilization capacity relative to other plant-associated strains in the primary phylogenetic logistic regression analysis (all phyloglm FDR q < 0.05; Fig. 5B). However, these family- and species-level associations were no longer significant when each focal group was compared only with other monocot-associated strains (all phyloglm FDR q > 0.05; Fig. S7). Thus, the apparent associations with individual monocot families and species largely reflected the broader depletion of disaccharide utilization among monocot-associated strains rather than additional depletion specific to those host taxa. Pagel correlated-evolution analyses showed a similar overall pattern (Fig. S7).

Together, these analyses reveal a consistent depletion of disaccharide-utilization capacity among monocot-associated RSSC strains that persists across alternative phylogenetic analyses, in the experimentally phenotyped subset, and after adjustment for geographic origin. These results show that the two metabolic traits underlying the classical biovar system differ not only in their genetic and evolutionary origins but also in their relationships with host association.

## Discussion

The classical RSSC biovar system has been used for more than half a century to describe variation in hexitol and disaccharide utilization, yet the molecular and evolutionary basis of these phenotypes has remained incompletely resolved. Here, we identify the *smo* locus and FAD-dependent glucose dehydrogenase (FADGDH) as major determinants of hexitol and disaccharide utilization, respectively, and show that the two metabolic systems have followed fundamentally different evolutionary trajectories. An ancestral *smo*-like system appears to have been lost early in RSSC evolution and subsequently reacquired in the ancestor of phylotype I through horizontal transfer from a *Burkholderia*-related lineage, whereas FADGDH was inherited vertically and recurrently lost in multiple RSSC lineages. Analyses across 1,082 strains further revealed a phylogenetically robust depletion of disaccharide-utilization capacity among monocot-associated strains. Together, these findings connect the molecular basis of classical biovar traits to broader processes of metabolic gene gain, loss, and ecological differentiation across RSSC.

An earlier comparative genomic study identified a 22-kb genomic island uniquely present in hexitol-positive RSSC strains and proposed that this region might contribute to hexitol utilization ^23^. By combining an expanded genomic dataset with machine-learning and genome-wide association analyses, followed by targeted gene-deletion experiments, we established the *smo* locus within this 22-kb region as a key determinant of hexitol utilization. In particular, the *smoEFGK* genes encode an ABC transporter for hexitol uptake, while neighboring genes encode enzymes involved in their subsequent metabolism (Fig. 3A). Deletion of *smoE* abolished utilization of all three tested hexitols, supporting a common requirement for the transporter, whereas deletion of *smoS* eliminated sorbitol and dulcitol utilization but retained mannitol utilization (Fig. 2G). This phenotype suggests a functional partitioning among the metabolic enzymes encoded within the locus, with SmoS contributing specifically to sorbitol and dulcitol metabolism while mannitol can be processed through a distinct enzymatic route, potentially involving MtlK. Thus, the *smo* locus provides an integrated transport-and-metabolism module underlying utilization of multiple related hexitol substrates.

In contrast to hexitol utilization, whose candidate genomic region had been identified previously, little was known about the genes responsible for disaccharide utilization in RSSC. Our analyses identify FADGDH as a key determinant of this phenotype. Deletion of the gene encoding the FADGDH α-subunit abolished utilization of all three disaccharides under the biovar assay conditions, demonstrating a requirement for FADGDH in GMI1000. FADGDH is a membrane-bound periplasmic dehydrogenase ^28^, and the homolog from *B. cepacia* (BcGDH) can directly oxidize disaccharides, including maltose ^26^. Conservation of the catalytic residues and the modeled substrate-binding architecture in RSSC FADGDH suggest a similar mechanism (Figure 4A, B). Together, these observations support the longstanding hypothesis that positive disaccharide reactions in RSSC biovar assays may result from direct oxidation of reducing sugars to form aldobionic acids rather than their hydrolysis and subsequent assimilation ^20^. Such a mechanism could also explain why GMI1000 can show a positive cellobiose reaction despite being unable to use cellobiose efficiently as a sole carbon source ^29^.

FADGDH presence alone, however, does not fully determine the disaccharide-utilization phenotype. In the 250-strain dataset, 42 of the 195 strains (∼21.5%) carrying FADGDH were recorded as unable to utilize the three disaccharides (Fig. 2F). This suggests that the absence of FADGDH is strongly associated with loss of the phenotype, whereas its presence is not sufficient to guarantee a positive reaction. Previous work has shown substantial variation in substrate utilization among RSSC strains despite highly conserved metabolic networks ^30^. In addition, studies of BcGDH have shown that proper folding and periplasmic localization of the catalytic α-subunit depend on the γ-subunit, while amino-acid substitutions in the α-subunit can alter substrate specificity ^26,28^. Thus, variation in FADGDH expression, maturation, or coding sequence could contribute to phenotypic differences among FADGDH-positive strains, none of which is captured by simple KO presence-absence annotation. Additional strain-specific factors are therefore needed to explain why some FADGDH-positive strains remain disaccharide-negative.

The evolutionary histories of the two systems illustrate how bacterial metabolic repertoires can be remodeled through different genomic processes. RSSC genomes are known to be highly dynamic, with extensive gene-content variation and horizontal acquisition documented across strains ^13,31^. The phylotype-I *smo* locus represents a clear case of gain in metabolic capacity through HGT, an important force driving bacterial evolution ^32^. Homologous *smo*-like loci occur in non-RSSC *Ralstonia*, but the phylotype-I *smo* locus instead groups phylogenetically with loci from *Burkholderia*-related bacteria (Fig. 3F). Its evolutionary distribution is therefore most consistent with loss of an ancestral *smo*-like system early in RSSC evolution, followed by reacquisition of a homologous metabolic module in the ancestor of phylotype I through HGT from a *Burkholderia*-related lineage. More broadly, this loss-and-reacquisition history shows how HGT can restore metabolic functions that were previously lost from an ancestral lineage, potentially allowing descendant lineages to regain access to substrates encountered in newly occupied or changing ecological niches.

FADGDH shows the opposite pattern. Its phylogeny supports inheritance from a common *Ralstonia* ancestor, followed by recurrent degeneration and loss during RSSC diversification. The occurrence of distinct pseudogene structures in different parts of the RSSC phylogeny, including independently arising disruptive mutations that generate similar loss patterns (Fig. 4D, E), indicates that loss of this metabolic function has occurred repeatedly rather than through a single ancestral event. Gene loss is increasingly recognized as an important component of bacterial evolution, particularly when maintaining a function provides little benefit or incurs physiological costs under particular conditions ^33^. In RSSC, the contrasting histories of the *smo* locus and FADGDH therefore illustrate how metabolic diversity can arise both through gain or re-acquisition of metabolic functions and through recurrent elimination of ancestrally inherited ones. These are not simply opposite changes in gene content; they represent distinct evolutionary routes by which closely related bacterial lineages arrive at different metabolic capacities.

The evolutionary changes at these metabolic loci may also have ecological consequences for RSSC diversification, although their effects are not equally resolvable from comparative data. Hexitols are available in plant-associated environments occupied by RSSC: mannitol accumulates in infected tomato xylem and can support bacterial growth ^29^, whereas sorbitol has been detected in the tomato rhizosphere ^34^. Reacquisition of the *smo* locus may therefore have expanded the range of carbon sources available to phylotype I strains and increased their metabolic flexibility in host-associated environments. Consistent with this possibility, transfer of a large GMI1000 genomic region containing the *smo* locus into the phylotype IV strain Psi07 increased aggressiveness on tomato, although the transferred region also contained additional genes (e.g., type III effector genes) and the phenotype cannot be attributed specifically to hexitol metabolism ^35^. Importantly, because the *smo* locus is essentially fixed in phylotype I and absent from the other phylotypes (Fig. 3B), hexitol-utilization capacity is almost completely confounded with deep phylogenetic lineage. Thus, the loss of statistical significance after phylogenetic correction does not exclude an ecological contribution of the *smo* system, but rather reflects the limited ability of the present comparative data to disentangle its effect from other phylotype I characteristics. Overall, the availability of hexitols in RSSC-associated environments and the near universal conservation of the locus in phylotype I are consistent with the possibility that *smo* reacquisition contributed to the metabolic and ecological diversification of this lineage.

The ecological pattern associated with disaccharide utilization is more distinctive. Cellobiose can be generated during plant cell-wall degradation, including through the activity of the RSSC cellobiohydrolase CbhA during host colonization ^36^, and maltose has been detected in tomato root exudates and among rhizosphere metabolite pools ^34,37^. These observations indicate that FADGDH substrates are available in RSSC living environments and suggest that their utilization may influence the ecological fitness of RSSC. At the same time, our comparative analysis revealed a consistent depletion of disaccharide-utilization capacity among monocot-associated strains. This association persisted under alternative treatments of phylogenetic relatedness, within the subset of strains with experimentally documented phenotypes, and after adjustment for geographic origin. One possible explanation is that the fitness value of maintaining FADGDH-mediated disaccharide oxidation differs among host-associated environments. If these substrates are less available, less useful, or otherwise provide insufficient benefit in particular niches, recurrent loss of the pathway could become selectively tolerable or even advantageous by eliminating the costs of maintaining an unnecessary metabolic system. Conversely, FADGDH could be preferentially retained where oxidation of reducing disaccharides provides a physiological or competitive benefit.

Overall, our study resolves the genetic basis of the classical RSSC biovar traits and shows how contrasting histories of gene acquisition and loss have generated metabolic diversity across the species complex. The association between disaccharide utilization and host affiliation is consistent with correlated metabolic and ecological evolution, although it does not establish a causal relationship between metabolic gene loss and host association. These findings illustrate how dynamic changes in bacterial gene repertoires can generate lineage-associated metabolic diversity with potential ecological consequences.

## Materials and Methods

### Dataset assembly and genomic feature matrix

Biovar information of 250 RSSC strains was collected from 58 published studies, the NCPPB (National Collection of Plant Pathogenic Bacteria) database, NCBI (National Center for Biotechnology Information) sample metadata, and our own experimental data (Table S1). Biovar assignments were converted into two binary metabolic traits according to the classical classification scheme (Table 1). Genome assemblies were downloaded from NCBI or generated from our own sequencing data. Protein sequences from the 250 RSSC genome assemblies were functionally annotated using KofamScan v1.3.0 with the KOfam database ^38^. A custom Python script was then used to count the number of proteins assigned to each KO identifier in each genome and to construct a feature matrix, with genomes as rows and KO identifiers as columns. In total, 2,566 distinct KO identifiers were identified across the 250 genomes included in the training dataset (Table S2). KO copy-number profiles were standardized to zero mean and unit variance before PCA. The association between hexitol and disaccharide utilization phenotypes was assessed using Pearson’s chi-square test with Yates’ continuity correction. The genome assemblies of the 1,082 RSSC strains used for the FADGDH loss analysis and host association analysis were obtained from the NCBI RefSeq database. Associated metadata, including isolation source, host species, and geographic information, were collected from the corresponding NCBI records (Table S5).

### Machine learning model construction and evaluation

Two tree-based machine learning models were constructed using XGBoost (v2.1.0-dev) ^39^. For each of hexitol-utilization and disaccharide-utilization phenotypes, both XGBClassifier and XGBRFClassifier were evaluated and the model with the best overall performance was selected for the final analysis. Finally, the hexitol utilization model was built using the random forest classifier XGBRFClassifier and the disaccharide utilization model was built using the gradient boosting tree classifier XGBClassifier. Predictions from the hexitol and disaccharide utilization models were subsequently integrated to generate the final biovar predictions. Default parameter settings were used for both models.

Model performance was assessed using repeated stratified cross-validation implemented with RepeatedStratifiedKFold in sklearn.model_selection v1.9.0, with 10 folds and three repeats (n_splits = 10, n_repeats = 3). Accuracy, balanced accuracy, precision, recall, F1 score, specificity, Matthews correlation coefficient (MCC), and Cohen’s kappa were calculated separately for each test fold and averaged across the 30 folds. For the multiclass biovar model, accuracy was calculated as the overall proportion of correctly classified samples; balanced accuracy, precision, recall, F1 score, and specificity were macro-averaged across the four classes; and MCC and Cohen’s kappa were calculated as overall multiclass measures. Accuracy and balanced accuracy are reported as mean ± standard deviation.

Receiver operating characteristic (ROC) curves, precision-recall curves, and confusion matrices were generated using stratified 10-fold cross-validation implemented with RepeatedStratifiedKFold in sklearn.model_selection (n_splits = 10, n_repeats = 1). For ROC analysis, ROC curves were calculated separately for each test fold from the predicted probabilities. Individual ROC curves were linearly interpolated onto a common false-positive-rate grid before averaging to obtain the mean ROC curve and corresponding mean area under the ROC curve (AUC). For the multiclass biovar model, class labels were binarized and a micro-average ROC curve was calculated using a one-vs-rest representation. Precision–recall curves were generated from pooled out-of-fold predicted probabilities across all cross-validation folds. For the binary hexitol and disaccharide utilization models, a single precision–recall curve was calculated for each phenotype. For the multiclass biovar model, precision–recall curves were calculated separately for each biovar class using a one-vs-rest strategy, and average precision (AP) was calculated for each curve. Confusion matrices were constructed by pooling the predicted and true labels from all test folds.

### Identification of phenotype-associated genomic features

After training the hexitol and disaccharide utilization models on the complete training dataset, the contributions of KEGG Orthology features to model predictions were interpreted using SHapley Additive exPlanations (SHAP). SHAP values were calculated using TreeExplainer in the SHAP Python package v0.52.0, with feature_perturbation=“tree_path_dependent” and model_output=“raw”. Feature importance was quantified as the mean absolute SHAP value across all strains, and features were ranked separately for the hexitol- and disaccharide-utilization models.

Genome-wide association analyses were performed using pyseer (v1.4.2) ^40^ to identify candidate KO features associated with hexitol or disaccharide utilization phenotypes. Population structure was accounted for using phylogenetic distances of 250 RSSC strains, which were subjected to multidimensional-scaling (MDS). The first three MDS components were included as covariates in the association models. Association statistics were calculated for each KO feature, and features were ranked according to their statistical association with the corresponding phenotype. The highest-ranking GWAS features were compared with the highest-ranking SHAP features to identify candidates supported independently by phenotype association and model interpretation.

### Construction of gene deletion mutants

Deletion mutants of *smoE* (RS_RS10750), *smoS* (RS_RS10765), and the gene encoding the FADGDH α-subunit (RS_RS03880) were constructed in GMI1000 following the methods described previously ^41^. DNA fragments of approximately 500 bp upstream and downstream of each target gene were PCR-amplified from GMI1000 genomic DNA using Phanta Max Super-Fidelity DNA Polymerase. The gentamicin resistance cassette was amplified from the pK18mobsacB plasmid. Primers used for mutant construction and verification are listed in Supplementary Table S9.

The upstream homologous arm, gentamicin resistance cassette, and downstream homologous arm were fused by overlap PCR to generate a linear deletion construct in which the target gene was replaced by the gentamicin resistance cassette. After purification, the fusion fragment was mixed with naturally competent GMI1000 cells and incubated on nitrocellulose membranes placed on CTG agar plates to facilitate natural transformation and homologous recombination. Transformants were selected on TTC agar plates supplemented with rifampicin and gentamicin. Candidate mutants were initially screened by PCR using primers specific to the gentamicin resistance cassette. Positive transformants were further verified by PCR using primers flanking the recombination region, followed by Sanger sequencing.

### Biovar assay

The utilization of three hexitols (mannitol, sorbitol, and dulcitol) and three disaccharides (lactose, maltose, and cellobiose) was determined using a pH-indicator assay based on acid production during carbohydrate metabolism. The standard basal medium for biovar determination was prepared by dissolving 1.0 g of NH₄H₂PO₄, 0.2 g of KCl, 0.2 g of MgSO₄·7H₂O, 1.0 g of Bacto peptone, and 80 mg of bromothymol blue in distilled water to a final volume of 1 L ^20^. The pH was adjusted to 7.4 with NaOH, after which 3.0 g of agar was added and the medium was autoclaved.

Stock solutions containing 10% (w/v) mannitol, sorbitol, dulcitol, lactose, maltose, cellobiose, or glucose were prepared, with glucose used as a positive substrate control. Mannitol and dulcitol solutions were gently heated to facilitate dissolution. The carbohydrate solutions were sterilized by filtration through 0.22-μm membrane filters and added to the biovar medium at a final concentration of 1% (w/v).

Bacterial strains were grown in TTC broth, harvested by centrifugation, resuspended in sterile distilled water, and adjusted to an OD₆₀₀ of 1.0. For each assay, 1 mL of medium containing 900 μL of basal medium and 100 μL of the corresponding carbohydrate solution was dispensed into a 2-mL microcentrifuge tube and inoculated with 20 μL of bacterial suspension. The cultures were incubated at 28°C for 10 days. Bromothymol blue has a pH transition range of 6.0–7.4, appearing blue at pH 7.4 and yellow below pH 6.0. Carbohydrate utilization was considered positive when the bromothymol blue indicator changed from green to yellow, indicating acid production during carbohydrate metabolism. Uninoculated biovar medium containing the corresponding carbohydrate was included as a negative control. Strains of biovar 3 (GMI1000) and biovar 4 (DHD47) were included as reference controls, with biovar 3 serving as a positive strain control. All assays were performed with three biological replicates.

### Phylogenetic analysis

To reconstruct the phylogenetic tree of the 250 strains in the training dataset, single-copy orthologous genes were identified using OrthoFinder v3.0.1b1 with default parameters ^42^. The amino acid sequences of these genes were aligned using MAFFT v7.526 ^43^, trimmed using trimAl v1.5.1 ^44^ and concatenated into a supermatrix using AMAS v1.0 ^45^. A maximum-likelihood phylogenetic tree was inferred using IQ-TREE v3.0.1 ^46^ with the best-fit substitution model selected using ModelFinder and branch support assessed using 1,000 ultrafast bootstrap replicates. The tree was visualized using iTOL ^47^. The same alignment, trimming, tree-inference, and visualization procedures were used in the subsequent phylogenetic analyses.

The evolutionary origin of the *smo* gene cluster was investigated using the protein sequences from strain GMI1000 (*smoS*, *smoE*, *smoF*, *smoG*, *smoK*, *mtlK*, *smoC*, *tagD*, and *tagE*) as BLASTP queries against bacterial reference genomes in the NCBI RefSeq database. An E-value cutoff of 1 × 10⁻⁵ was used. Protein sequences from 22,260 bacterial genome assemblies representing distinct species were used to construct the BLAST database. The top 1,000 homologs of each protein were retained for individual gene tree reconstruction. Taxa that clustered with RSSC in these gene trees were selected for subsequent gene cluster analysis. Homologous gene clusters in the selected genomes were identified using cblaster v1.4.0 ^48^. Protein sequences of seven genes present in most of the selected taxa (*smoS*, *smoE*, *smoF*, *smoG*, *smoK*, *mtlK*, and *smoC*) were concatenated to reconstruct the gene cluster phylogeny. The *FADGDH* gene cluster phylogeny was reconstructed from concatenated amino acid sequences of the three subunit genes, represented by the GMI1000 locus tags *RS_RS03875* (γ subunit), *RS_RS03880* (α subunit), and *RS_RS03885* (β subunit), using the same procedure described above.

To test whether the placement of RSSC *smo* homologs with *Burkholderia*-related sequences was significantly better supported than alternative relationships with non-RSSC *Ralstonia*, constrained phylogenies were constructed in which the RSSC sequences were forced to cluster with homologs from non-RSSC *Ralstonia* species. The unconstrained and constrained topologies were compared in IQ-TREE ^46^ using the bootstrap proportion calculated by the resampling estimated log-likelihood method (bp-RELL), the Shimodaira–Hasegawa (SH) test, and the approximately unbiased (AU) test, based on 10,000 RELL replicates. The AU and SH tests return P values, whereas bp-RELL returns posterior weights. The AU test was used as the primary significance test, with constrained topologies rejected at P < 0.05. SH test P values and bp-RELL support weights were used as additional evidence.

The phylogeny of 1,082 RSSC strains was reconstructed using single copy BUSCO genes identified by BUSCO v6.0.0 ^49^ with the *ralstonia*_odb12 lineage dataset. A total of 277 genes present as single-copy orthologs in all genomes were concatenated for phylogenetic reconstruction.

### Protein structure analysis

The crystal structure of BcGDH (PDB ID: 8GRJ; chains A and B) was used as the template to construct a structural model of the RsGDH αγ complex using SWISS-MODEL ^50^. The ligand structures of glucose, lactose, maltose, and cellobiose were obtained from PubChem under CID entries 5793, 440995, 439186, and 439178, respectively.

Glucose was docked into the RsGDH αγ model without positional constraints using Schrödinger Maestro (Release 2023-1). Ligand preparation, protein preparation, and ligand docking were performed using the corresponding workflows in Maestro, with docking carried out using Glide in Extra Precision (XP) mode. Among the resulting poses, the pose most consistent with the ligand binding mode observed in the BcGDH crystal structure was selected for subsequent structural analysis. For comparison of ligand binding poses, the RsGDH model and the BcGDH crystal structure were superposed using the “align” command in PyMOL v3.1.6.1, and the resulting structural superposition was visualized in PyMOL.

The binding poses of lactose, maltose, and cellobiose were generated by aligning the C1–C6 atoms of the glucose moiety at the reducing end of each disaccharide with the corresponding atoms of D-glucono-1,5-lactone in the BcGDH crystal structure using the “pair_fit” command in PyMOL. The resulting protein–ligand complexes were processed using the protein preparation workflow in Schrödinger and subsequently energy minimized using MacroModel with the OPLS4 force field and no solvent model.

### Gene cluster analysis

All gene clusters analyzed in this study were identified using cblaster v1.4.0 ^48^. Local cblaster databases were constructed from genome annotation files in GenBank format. Default filtering thresholds were used: --min_hits 3, --unique 3, --gap 20000, --min_identity 30, --min_coverage 50, --max_evalue 0.01. Detailed results are provided in Tables S10–S14.

For the *smo* locus, the corresponding GMI1000 protein sequences were used as query sequences. Locus presence was determined in the 250-strain dataset (Table S10). In the 1,082-strain dataset, locus structure was further classified according to the presence or absence of individual *smo* locus genes (Table S12, S15). Gene cluster structures displayed alongside the *smo* gene cluster phylogeny were visualized using clinker v0.0.32 ^51^.

For the *FADGDH* locus, the three FADGDH subunit genes from RSSC strain HA4-1 (GCA_003999715.1), together with six conserved flanking genes (three upstream and three downstream), were used as query sequences. Inclusion of the flanking genes allowed the corresponding genomic region to be identified even when one or more FADGDH subunit genes were absent or annotated as pseudogenes. The *FADGDH* locus was defined by the positions of the conserved flanking genes. The presence of each FADGDH subunit gene was determined from cblaster hits, whereas pseudogene annotations were obtained from the corresponding GenBank files. The genomic context of the locus was visualized using the pyGenomeViz python package 1.6.1, and structural types were classified accordingly (Fig. S5; Table S13). For subunits annotated as pseudogenes, the annotated sequences were extracted and aligned to the corresponding GMI1000 reference region using MAFFT ^43^. The resulting alignments were examined using a custom Python script to classify pseudogenization patterns.

### Association analyses of metabolic phenotypes with hosts and isolation source

Host, isolation source, and geographic location metadata for the 1,082 RSSC strains were collected from NCBI metadata. Strains with reported isolation-source and host information were assigned to one of three broad categories (plant, soil/rhizosphere, or water). Plant-associated strains were further classified according to host clade (monocotyledon or dicotyledon), host family, and host species. Of the 1,082 strains, 1,035 had an assigned broad isolation source, and 852 plant-associated strains had host clade, family, and species assignments. Comparisons were performed separately for isolation source, host clade, host family, and host species. For isolation source analyses, each focal source was compared with all other sources. For host clade, host family, and host species analyses, each focal group was compared with all other plant-associated strains. Hexitol- and disaccharide-utilization phenotypes were predicted using the machine learning models for the 858 strains without experimentally documented biovar phenotypes. Detailed metadata are provided in Table S5.

Associations between metabolic phenotypes and host or isolation source categories were primarily tested using Ives–Garland phylogenetic logistic regression ^52^, implemented using the phyloglm function in the phylolm R package v2.6.5 ^53^. Zero length branches in the 1,082-strain phylogeny were assigned a small positive length of 1× 10⁻¹⁰ for the analyses. Effect sizes were expressed as odds ratios (ORs), with OR > 1 indicating enrichment and OR < 1 indicating depletion of the corresponding metabolic phenotype in the focal group. P values were adjusted for multiple testing using the false discovery rate (FDR), with FDR q < 0.05 considered statistically significant.

Several sensitivity analyses were performed. First, phylogenetic structure was alternatively represented by principal-coordinate/multidimensional-scaling (PC/MDS) axes. For each focal-versus-reference comparison, pairwise patristic distances among all included strains were calculated from the corresponding pruned phylogeny. Classical multidimensional scaling was then performed on the distance matrix, and the first five principal coordinate axes were included as covariates in a binomial logistic regression model. Firth penalized logistic regression was used when separation was detected. Second, Bayesian phylogenetic generalized linear mixed models were fitted using the brms R package v2.23.0 ^54^. Disaccharide utilization capacity was treated as a binary response, with focal group membership included as a fixed effect and phylogenetic relationships among strains incorporated as a random effect structured by a phylogenetic correlation matrix derived from the corresponding pruned tree. Models were fitted using 10 chains of 2,000 iterations, including 1,000 warmup iterations. Posterior mean odds ratios were used to determine the direction of the focal group effect. In addition, to assess whether geographic distribution contributed to the observed associations, continent was included as a categorical covariate in the phyloglm model. An experimental-only analysis was performed using strains with experimentally supported sugar utilization phenotypes. To determine whether host family and species associations reflected the broader monocot/dicot pattern, additional analyses compared each focal host family or species only with other hosts of the same cotyledon type.

Correlated evolution between ecological association and disaccharide utilization capacity was tested using Pagel’s discrete method implemented with the fitMk function in the phytools R package v2.5.2 ^55^. The independent model comprised two separate equal rates (ER) models, whereas the dependent model was fitted as a four-state all-rates-different (ARD) model. The independent and dependent models were compared using a parametric bootstrap likelihood ratio test based on 500 replicates simulated under the independent model, with bootstrap P < 0.05 considered significant.

## Resource availability

### Lead contact

Requests for further information and resources should be directed to and will be fulfilled by the lead contact, Xiaofan Zhou.

### Materials availability

Bacterial strains generated in this study are available from the lead contact upon reasonable request.

### Data and code availability

Genome assemblies analyzed in this study are publicly available from NCBI, and the corresponding accession numbers are provided in Table S1. Genome data for strains collected by our laboratory are available from the lead contact upon reasonable request. The analysis code used in this study is available from the lead contact upon reasonable request.

## Acknowledgements

We thank Shenjun Zhu and Jiancheng Luo for helpful suggestions on methodology. This work was supported by the National Natural Science Foundation of China (NSFC 32470184) and Guangdong S&T Program (2026B0202190005).

## Author contributions

Conceptualization, X.Z.; data curation, Y.L.; formal analysis, Y.L. and X.Z.; investigation, Y.L., S.Y., and Q.L.; methodology, Y.L. and X.Z.; supervision, X.Z.; visualization, Y.L.; writing – original draft, Y.L.; writing – review & editing, Y.L. and X.Z.

## Supplemental information

Document S1. Figures S1–S7 and Table S4 Data S1. Tables S1–S3 and S5–S15

---

**Figure S1.**
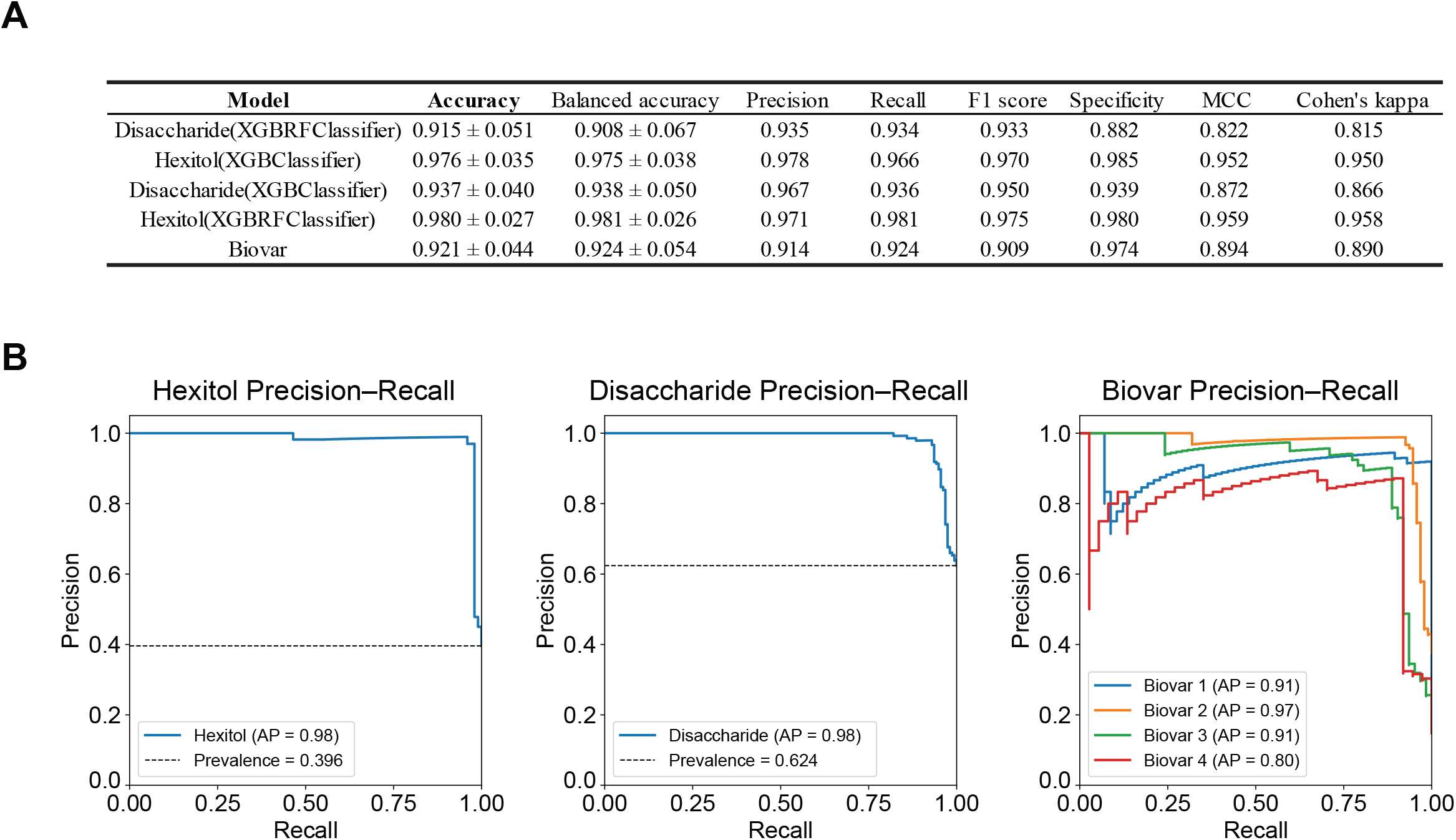
Performance evaluation of machine learning models. (A) Performance of the candidate and selected models for hexitol utilization, disaccharide utilization, as well as the integrated biovar model. Model performance was evaluated using repeated stratified crossvalidation. Accuracy and balanced accuracy are presented as the mean ± standard deviation. Additional performance metrics include precision, recall, F1 score, specificity, Matthews correlation coefficient (MCC), and Cohen’s kappa. (B) Precision-recall curves for the selected models for hexitol utilization, disaccharide utilization, and the four biovar classes. Biovar performance was evaluated using a one-vs-rest strategy. AP indicates average precision. Dashed horizontal lines indicate the prevalence of the positive class in binary classification tasks.

**Figure S2.**
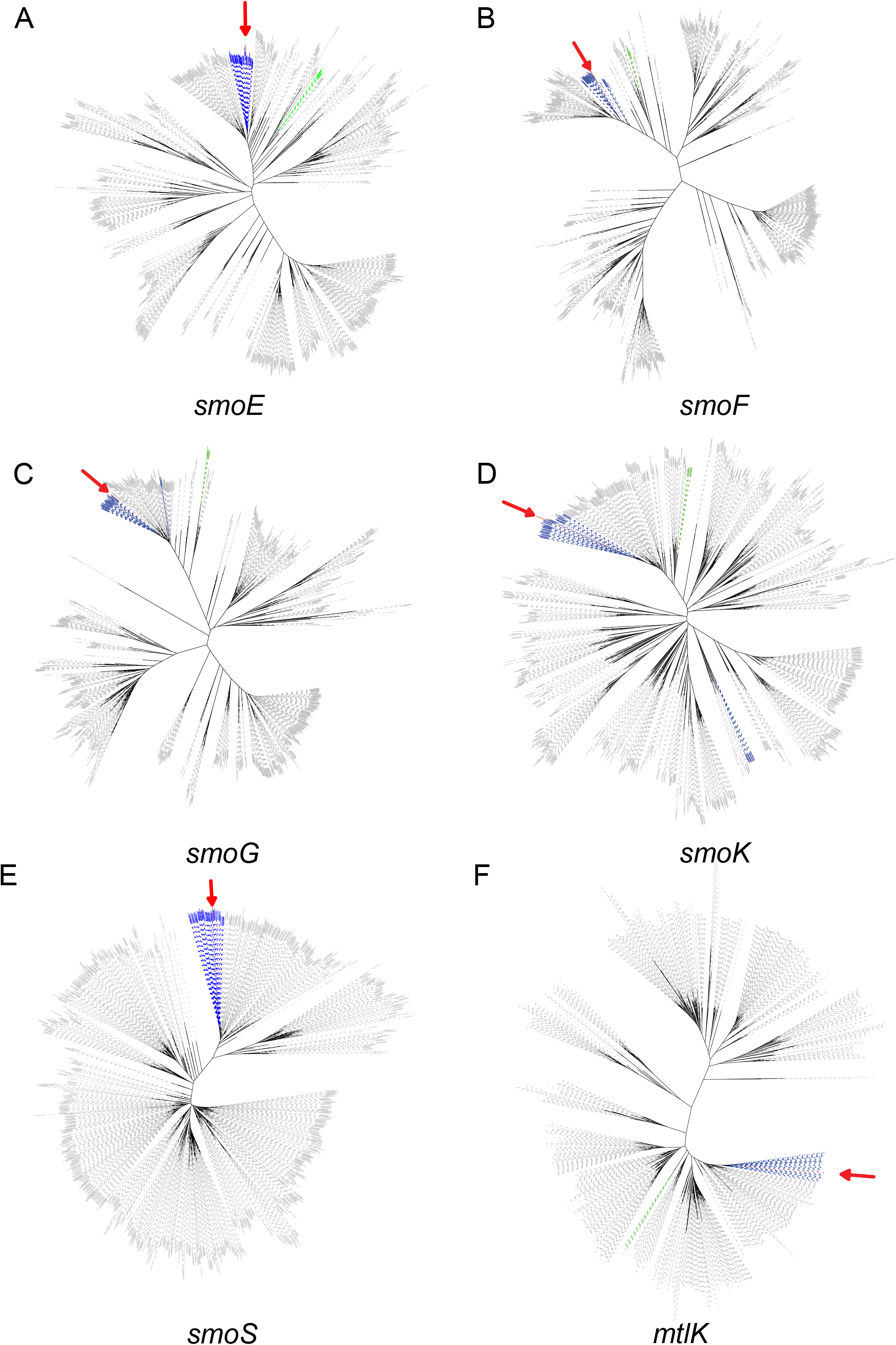
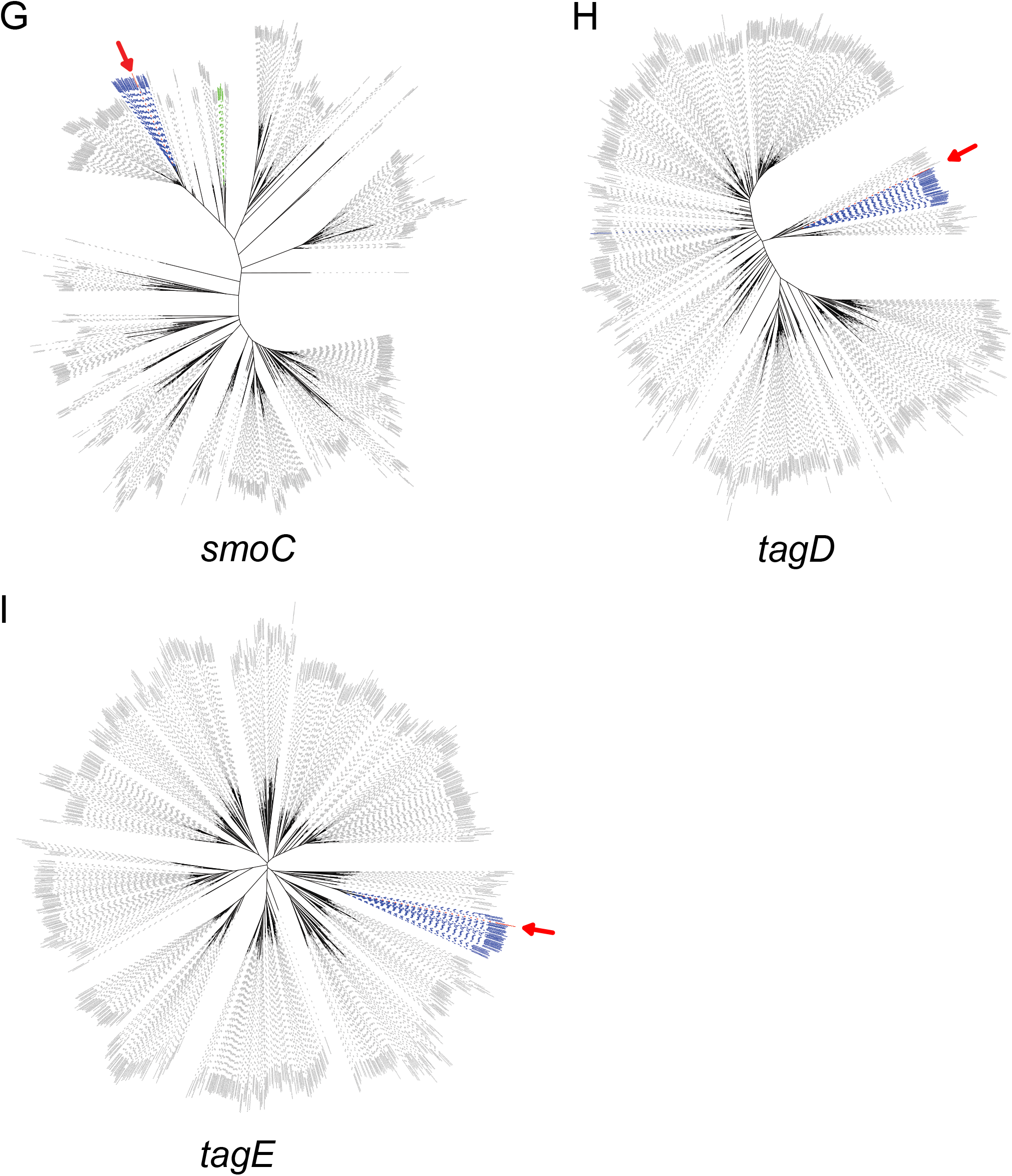
Phylogenies of individual genes in the GMI1000 *smoEFGK* gene cluster. RSSC is highlighted in red and indicated by a red arrow. Other Ralstonia species are colored green, and Burkholderia species are colored blue. Protein sequences from R. pseudosolanacearum GMI1000 (*smoE*, RS_RS10750; *smoF*, RS_RS10745; *smoG*, RS_RS10740; *smoK*, RS_RS10730; *smoS*, RS_RS10765; *mtlK*, RS_RS10685; *smoC*, RS_RS10675; *tagD*, RS_RS10760; *tagE*, RS_RS10755) were used as BLASTP queries to search for homologs in the NCBI RefSeq bacterial reference assemblies. The top 1,000 homologs for each protein were retained, aligned and trimmed. Maximum-likelihood trees were reconstructed using IQ-TREE with 1,000 ultrafast bootstrap replicates.

**Figure S3.**
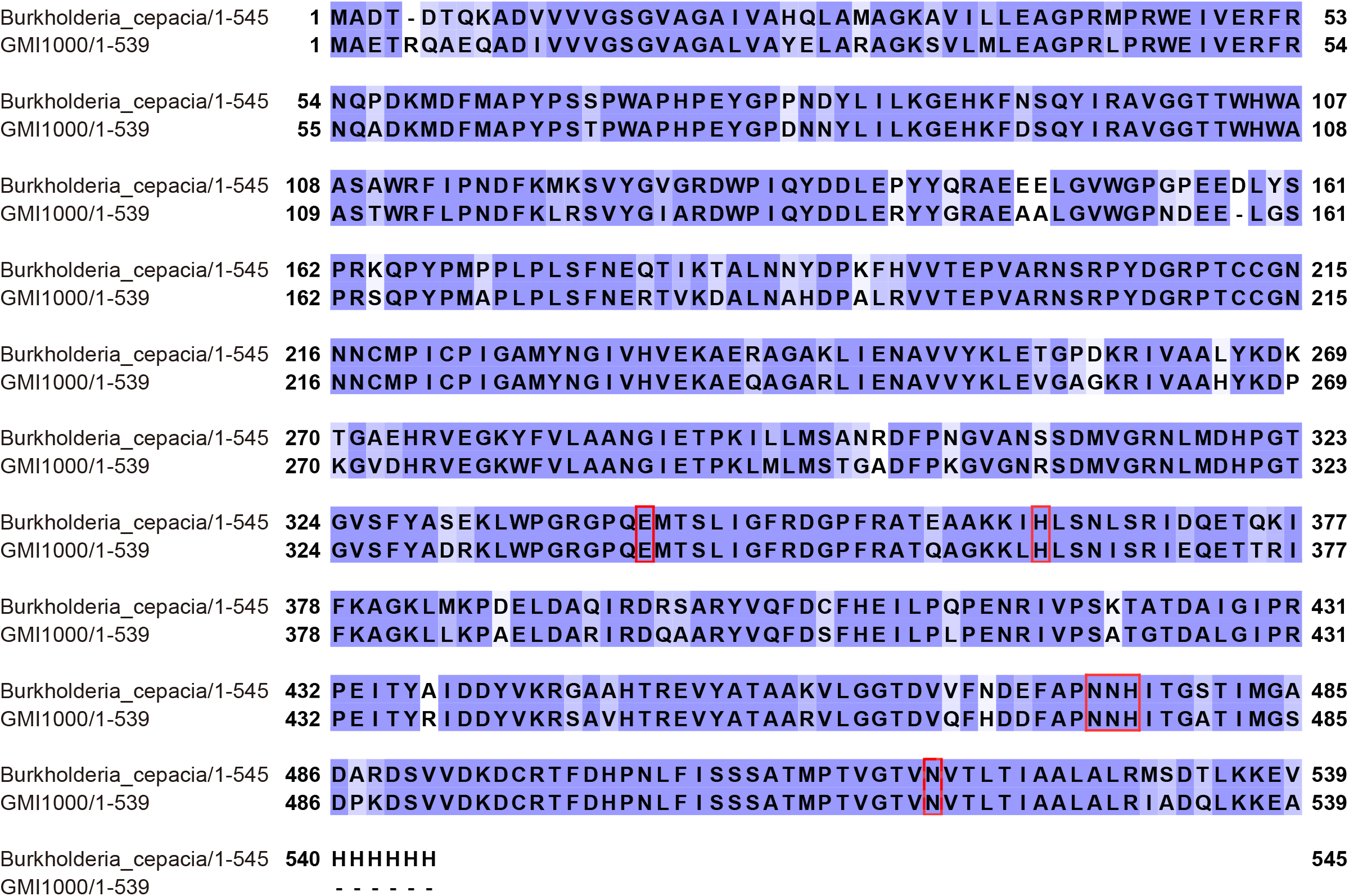
Protein sequence alignment of the FADGDH α-subunit from GMI1000 and *Burkholderia cepacia*. Active site residues identified in BcGDH are marked with red boxes and are identical between the two proteins.

**Figure S4.**
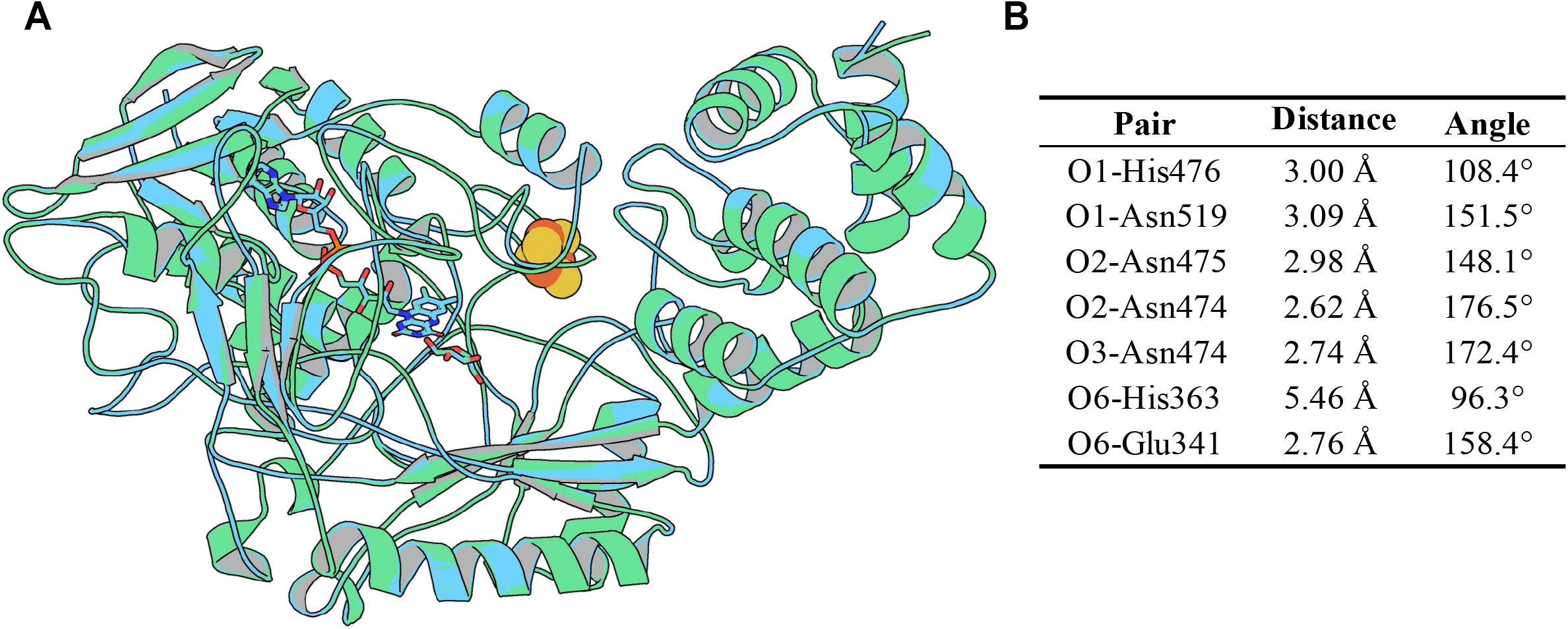
Structural superposition of RsGDH and BcGDH. (A)The homology modeled RsGDH structure (blue) was superimposed onto the BcGDH crystal structure (green). FAD and D-glucono-1,5-lactone (LGC) are shown as sticks. The iron-sulfur cluster is shown as spheres. (B) Assessment of hydrogen-bond geometry in the docked RsGDH-glucose complex for glucose-residue pairs corresponding to hydrogen bonds observed in the BcGDH crystal structure.

**Figure S5.**
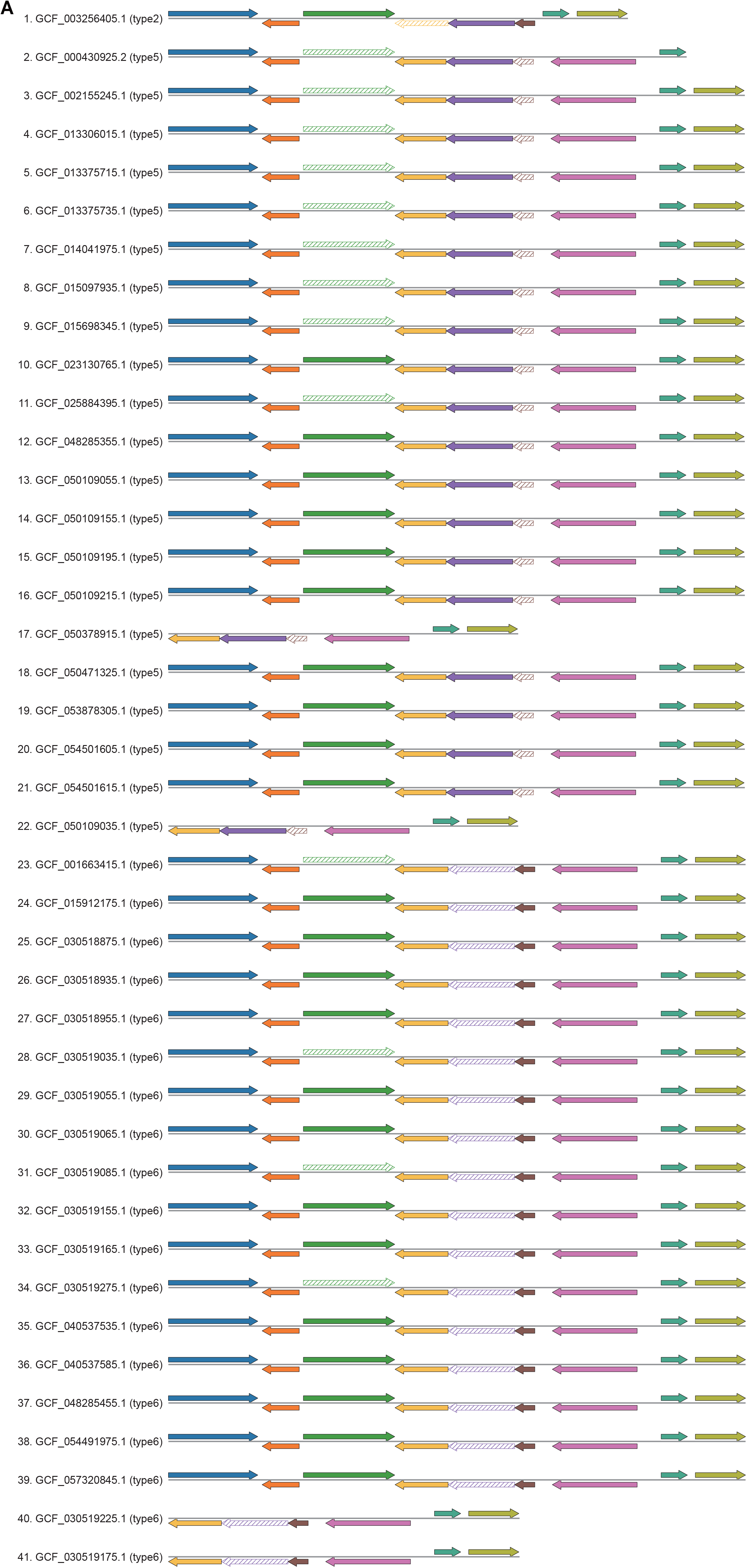
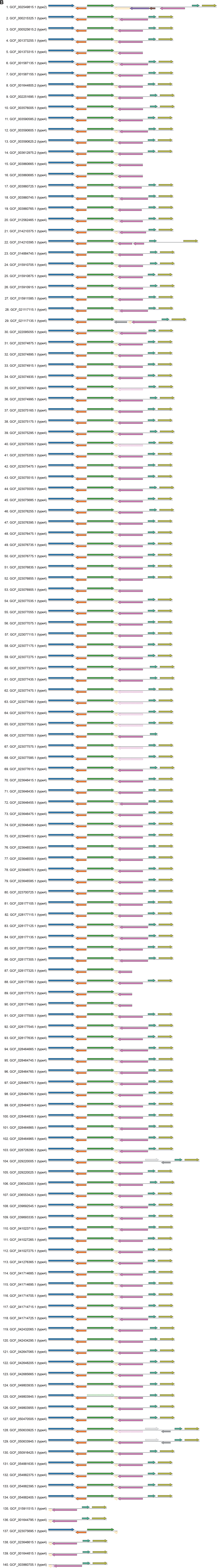
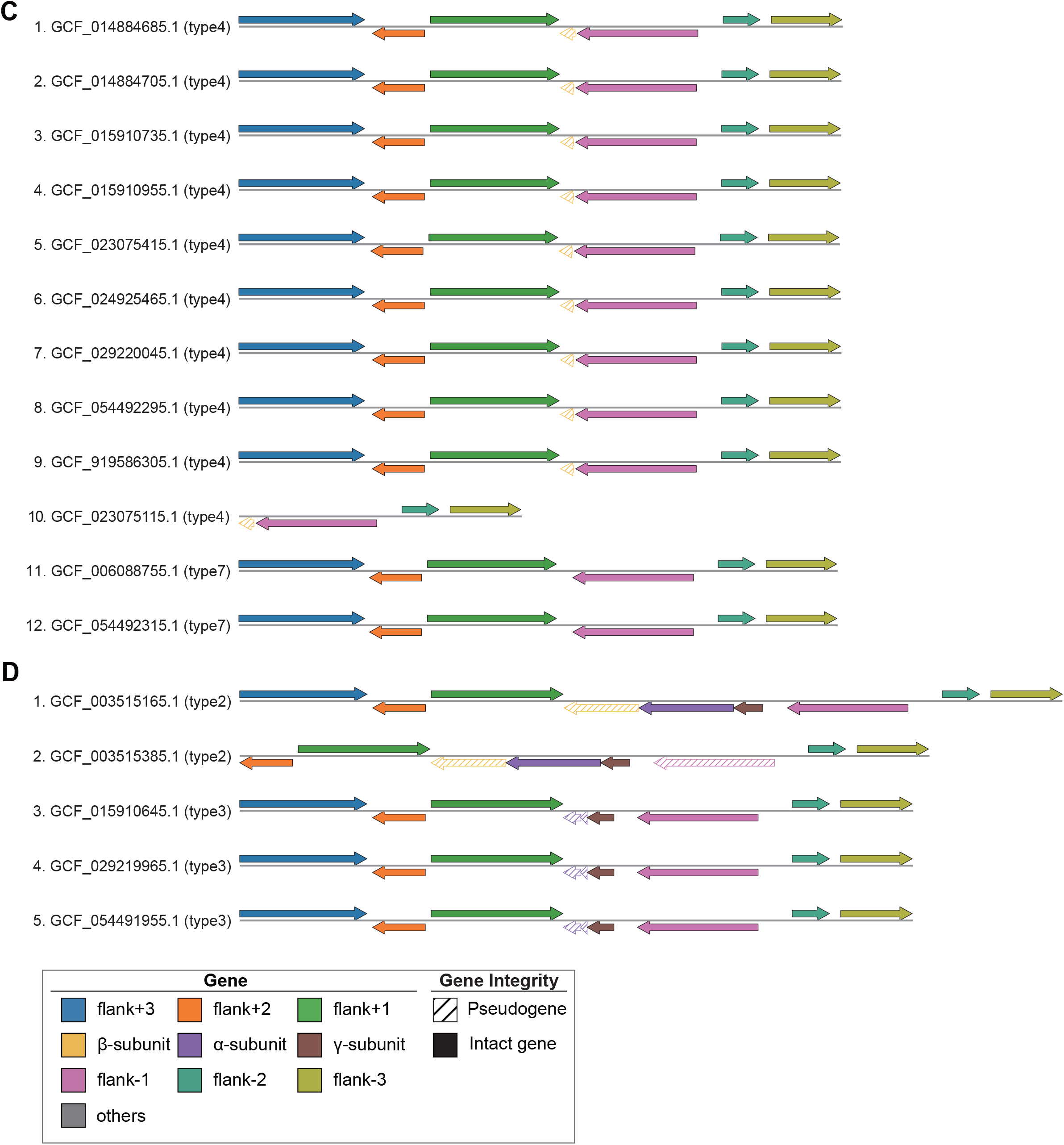
FADGDH locus structures associated with degeneration in RSSC genomes. Gene cluster structures are grouped by phylotype: phylotype I (A), phylotype II (B), phylotype III (C), and phylotype IV (D). Conserved flanking genes are included. Genomes carrying the intact FADGDH gene cluster (type 1) are not shown. Pseudogenes are indicated by diagonally hatched arrows. Grey arrows indicate non-conserved genes inserted within the conserved flanking region.

**Figure S6.**
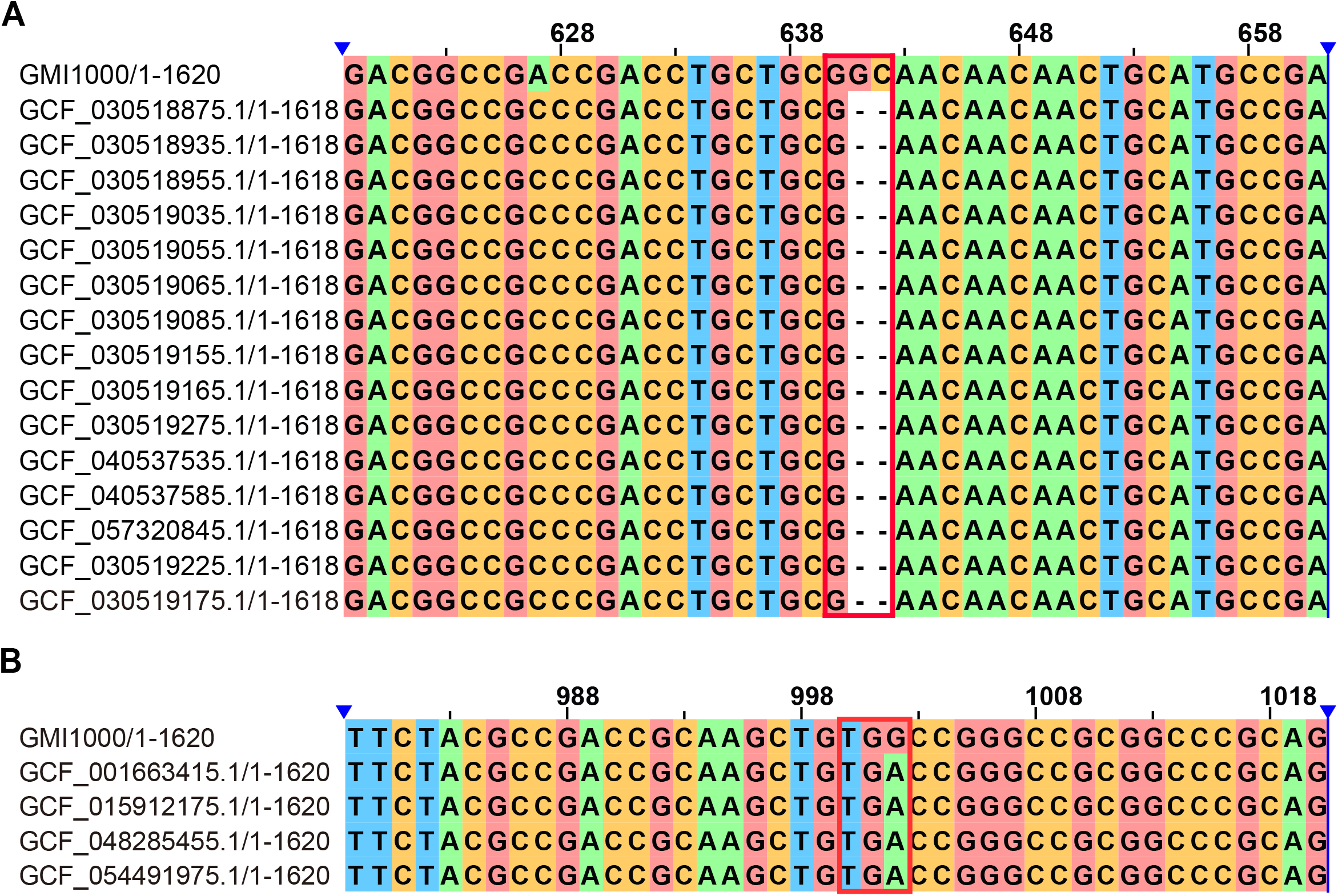
Nucleotide sequence alignments of the FADGDH α-subunit gene. Sequences carrying the type 6 frameshift mutation (A) and internal stop-codon mutation (B) were aligned to the intact GMI1000 sequence. Mutated sites are indicated by red boxes.

**Figure S7.**
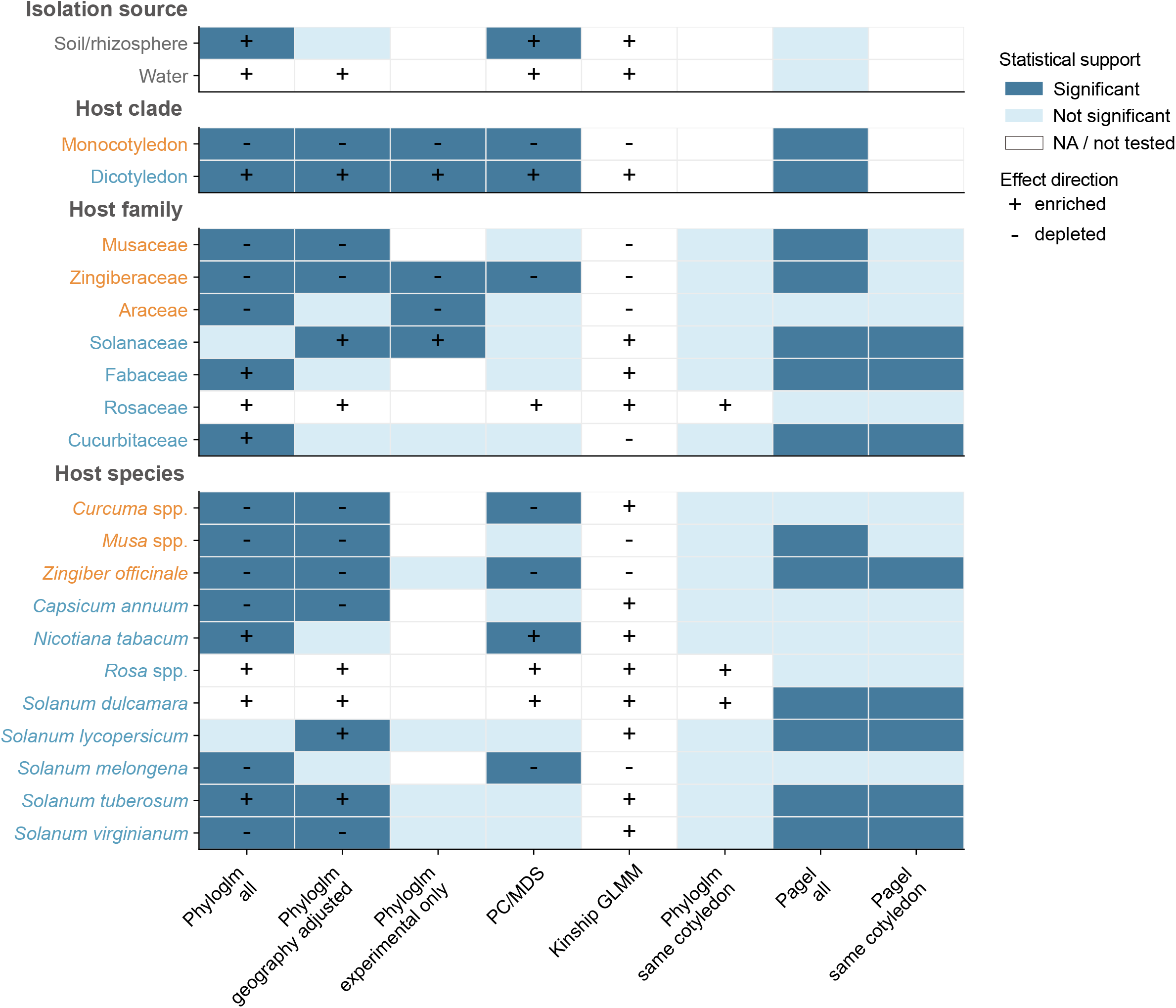
Sensitivity analysis summary of disaccharide utilization associations with host categories and isolation source. Dark blue cells indicate statistically significant support, light blue cells indicate non-significant results, and white cells indicate entries that were not included in the corresponding analysis or for which no P value or FDR-adjusted significance was available. Plus and minus signs denote enrichment and depletion of disaccharide utilization capacity in the focal group, respectively, based on the direction of the effect estimate from each analysis. The “Phyloglm all” column shows the primary Ives-Garland phylogenetic logistic regression results comparing each focal group with the corresponding full reference set: each isolation source was compared with all other sources, including plant associated strains; monocot associated strains were compared with dicot associated strains for host clade contrasts; host family and host species groups were compared with all other plant associated strains. The “Phyloglm geography adjusted” column shows the same phyloglm contrasts after adding geographic continent as an additional covariate. The “Phyloglm experimental only” column shows phyloglm results restricted to strains with experimentally supported utilization phenotypes. The “PC/MDS” column shows logistic models adjusted by phylogenetic ancestry axes derived from the RSSC phylogeny. The “Kinship GLMM” column shows only the estimated direction of the focal-group effect from a Bayesian phylogenetic GLMM that accounts for strain relatedness using a tree-derived covariance matrix; no significance color is assigned to this column. The “Phyloglm same cotyledon” column compares focal host family or host species groups only with other hosts of the same cotyledon type. The “Pagel all” column shows Pagel correlated evolution tests comparing dependent and independent evolution of the focal group and disaccharide utilization using the same full reference set as in “Phyloglm all”. The “Pagel same cotyledon” column shows the corresponding Pagel test using the same cotyledon comparison as in “Phyloglm same cotyledon”. Water, Rosaceae, *Rosa* spp., and *Solanum dulcamara* showed complete separation because all focal-group strains were disaccharide-positive; their direction was shown as enrichment, but standard adjusted OR and significance estimates were not interpreted in phyloglm-based and PC/MDS analyses. Statistical significance was assessed at FDR q < 0.05 for phyloglm and PC/MDS analyses, and at bootstrap P < 0.05 for Pagel correlated evolution analyses. Orange and blue labels indicate monocot and dicot hosts, respectively. Detailed values are provided in Tables S6-S8.

**Table S4.** Topology tests comparing two hypotheses for the origin of the *smoEFGK* geneclusterinRSSC. BO, *Burkholderia* origin;non-BO, *non-Burkholderia* origin; logL, log likelihood of the tested topology; deltaL, difference from the maximal log likelihood in the set; p-AU, p value of the approximately unbiased test; bp-RELL, bootstrap proportion calculated using RELL method; p-SH, p value of Shimodaira-Hasegawa test.

| **Gene** | **Hypothesis** | **logL** | **deltaL** | **p-AU** | **bp-RELL** | **p-SH** |
| --- | --- | --- | --- | --- | --- | --- |
| *smoE* | BO | -25441.3173 | 0.0000 | 0.9990 | 1.0000 | 1.0000 |
| *smoE* | Non-BO | -25624.0140 | 182.7000 | 0.0015* | 0.0003 | 0.0003* |
| *smoF* | BO | -15697.9309 | 0.0000 | 0.9990 | 0.9990 | 1.0000 |
| *smoF* | Non-BO | -15805.7019 | 107.7700 | 0.0011* | 0.0011 | 0.0014* |
| *smoG* | BO | -7348.2924 | 0.0000 | 0.9490 | 0.9410 | 1.0000 |
| *smoG* | Non-BO | -7404.5525 | 56.2600 | 0.0508 | 0.0589 | 0.0617 |
| *smoK* | BO | -20198.2560 | 0.0000 | 0.9790 | 0.9790 | 1.0000 |
| *smoK* | Non-BO | -20283.5368 | 85.2810 | 0.0206* | 0.0206 | 0.0183* |
| *smoC* | BO | -15219.1115 | 0.0000 | 1.0000 | 1.0000 | 1.0000 |
| *smoC* | Non-BO | -15366.7816 | 147.6700 | 4.56e-07* | 0.0000 | 0.0000* |
| *mtlK* | BO | -66668.5911 | 0.0000 | 0.9680 | 0.9650 | 1.0000 |
| *mtlK* | Non-BO | -66781.1279 | 112.5400 | 0.0316* | 0.0349 | 0.0322* |

## Supplementary Information Contents

**Figure S1.** Performance evaluation of machine learning models.

**Figure S2.** Phylogenies of individual genes in the GMI1000 *smoEFGK* gene cluster.

**Figure S3.** Protein sequence alignment of the FADGDH α-subunit from GMI1000 and *Burkholderia cepacia*.

**Figure S4.** Structural superposition of RsGDH and BcGDH.

**Figure S5.** FADGDH gene cluster structures associated with locus degeneration in RSSC genomes.

**Figure S6.** Nucleotide sequence alignments of the FADGDH α-subunit gene.

**Figure S7.** Sensitivity analysis summary of disaccharide utilization associations with host categories and isolation source.

**Table S4.** Topology tests comparing two hypotheses for the origin of the smoEFGK

## References

1. David, K.T., Schraiber, J.G., Crandall, J.G., Labella, A.L., Opulente, D.A., Harrison, M.-C., Wolters, J.F., Zhou, X., Shen, X.-X., Groenewald, M., et al. (2025). Convergent expansions of keystone gene families drive metabolic innovation in Saccharomycotina yeasts. Proc. Natl. Acad. Sci. U.S.A. 122, e2500165122. 10.1073/pnas.2500165122.

2. Näsvall, J., Sun, L., Roth, J.R., and Andersson, D.I. (2012). Real-time evolution of new genes by innovation, amplification, and divergence. Science 338, 384–387. 10.1126/science.1226521.

3. Slot, J.C., and Rokas, A. (2010). Multiple *GAL* pathway gene clusters evolved independently and by different mechanisms in fungi. Proc. Natl. Acad. Sci. U.S.A. 107, 10136–10141. 10.1073/pnas.0914418107.

4. D’Souza, G., Waschina, S., Pande, S., Bohl, K., Kaleta, C., and Kost, C. (2014). Less is more: Selective advantages can explain the prevalent loss of biosynthetic genes in bacteria: adaptive loss of biosynthetic genes in bacteria. Evolution 68, 2559–2570. 10.1111/evo.12468.

5. Genin, S., and Denny, T.P. (2012). Pathogenomics of the *Ralstonia solanacearum* species complex. Annu. Rev. Phytopathol. 50, 67–89. 10.1146/annurev-phyto-081211-173000.

6. Vailleau, F., and Genin, S. (2023). *Ralstonia solanacearum*: An arsenal of virulence strategies and prospects for resistance. Annu. Rev. Phytopathol. 61, 25–47. 10.1146/annurev-phyto-021622-104551.

7. Lowe-Power, T.M., Avalos, J., Bai, Y., Munoz, M.C., Chipman, K., Elmgreen, V.N., Prasad, N., Ramirez, B., Sandhar, A., Tom, C.E., et al. (2024). A meta-analysis of the known global distribution and host range of the *Ralstonia* species complex. Preprint at bioRxiv, 10.1101/2020.07.13.189936.

8. Barik, S., Reddy, A.C., Ponnam, N., Kumari, M., Acharya, G.C., Lakshmana Reddy, D.C., Petikam, S., and Sahu, G.S. (2020). Breeding for bacterial wilt resistance in eggplant (*Solanum melongena* L.): Progress and prospects. Crop Prot. 137, 105270. 10.1016/j.cropro.2020.105270.

9. Álvarez, B., López, M.M., and Biosca, E.G. (2008). Survival strategies and pathogenicity of *Ralstonia solanacearum* phylotype II subjected to prolonged starvation in environmental water microcosms. Microbiology 154, 3590–3598. 10.1099/mic.0.2008/019448-0.

10. van Elsas, J.D., Kastelein, P., De Vries, P.M., and Van Overbeek, L.S. (2001). Effects of ecological factors on the survival and physiology of *Ralstonia solanacearum* bv. 2 in irrigation water. Can. J. Microbiol. 47, 842–854. 10.1139/cjm-47-9-842.

11. van Elsas, J.D., Kastelein, P., Van Bekkum, P., Van Der Wolf, J.M., De Vries, P.M., and Van Overbeek, L.S. (2000). Survival of *Ralstonia solanacearum* biovar 2, the causative agent of potato brown rot, in field and microcosm soils in temperate climates. Phytopathology 90, 1358–1366. 10.1094/PHYTO.2000.90.12.1358.

12. Guidot, A., Coupat, B., Fall, S., Prior, P., and Bertolla, F. (2009). Horizontal gene transfer between *ralstonia solanacearum* strains detected by comparative genomic hybridization on microarrays. The ISME Journal 3, 549–562. 10.1038/ismej.2009.14.

13. Salanoubat, M., Genin, S., Artiguenave, F., Gouzy, J., Mangenot, S., Arlat, M., Billault, A., Brottier, P., Camus, J.C., Cattolico, L., et al. (2002). Genome sequence of the plant pathogen *Ralstonia solanacearum*. Nature 415, 497–502. 10.1038/415497a.

14. Lefeuvre, P., Cellier, G., Remenant, B., Chiroleu, F., and Prior, P. (2013). Constraints on genome dynamics revealed from gene distribution among the ralstonia solanacearum species. PLoS ONE 8, e63155. 10.1371/journal.pone.0063155.

15. Ahmed, W., Yang, J., Tan, Y., Munir, S., Liu, Q., Zhang, J., Ji, G., and Zhao, Z. (2022). *Ralstonia solanacearum*, a deadly pathogen: Revisiting the bacterial wilt biocontrol practices in tobacco and other solanaceae. Rhizosphere 21, 100479. 10.1016/j.rhisph.2022.100479.

16. de Pedro-Jové, R., Corral, J., Rocafort, M., Puigvert, M., Azam, F.L., Vandecaveye, A., Macho, A.P., Balsalobre, C., Coll, N.S., Orellano, E., et al. (2023). Gene expression changes throughout the life cycle allow a bacterial plant pathogen to persist in diverse environmental habitats. PLoS Pathog. 19, e1011888. 10.1371/journal.ppat.1011888.

17. Lowe-Power, T.M., Khokhani, D., and Allen, C. (2018). How *Ralstonia solanacearum* exploits and thrives in the flowing plant xylem environment. Trends Microbiol. 26, 929–942. 10.1016/j.tim.2018.06.002.

18. Peyraud, R., Cottret, L., Marmiesse, L., Gouzy, J., and Genin, S. (2016). A resource allocation trade-off between virulence and proliferation drives metabolic versatility in the plant pathogen *Ralstonia solanacearum*. PLoS Pathog. 12, e1005939. 10.1371/journal.ppat.1005939.

19. Riesco, R., and Trujillo, M.E. (2024). Update on the proposed minimal standards for the use of genome data for the taxonomy of prokaryotes. Int. J. Syst. Evol. Microbiol. 74, 006300. 10.1099/ijsem.0.006300.

20. Hayward, A.C. (1964). Characteristics of *Pseudomonas solanacearum*. J. Appl. Bacteriol. 27, 265–277. 10.1111/j.1365-2672.1964.tb04912.x.

21. Prior, P., and Fegan, M. (2005). Recent developments in the phylogeny and classification of *Ralstonia solanacearum*. Acta Hortic. 695, 127–136. 10.17660/ActaHortic.2005.695.14.

22. Safni, I., Cleenwerck, I., De Vos, P., Fegan, M., Sly, L., and Kappler, U. (2014). Polyphasic taxonomic revision of the *Ralstonia solanacearum* species complex: proposal to emend the descriptions of *Ralstonia solanacearum* and *Ralstonia syzygii* and reclassify current *R. syzygii* strains as *Ralstonia syzygii* subsp. *syzygii* subsp. nov., *R. solanacearum* phylotype IV strains as *Ralstonia syzygii* subsp. *indonesiensis* subsp. nov., banana blood disease bacterium strains as *Ralstonia syzygii* subsp. *celebesensis* subsp. nov. and *R. solanacearum* phylotype I and III strains as *Ralstonia pseudosolanacearum* sp. nov. Int. J. Syst. Evol. Microbiol. 64, 3087–3103. 10.1099/ijs.0.066712-0.

23. Gabriel, D.W., Allen, C., Schell, M., Denny, T.P., Greenberg, J.T., Duan, Y.P., Flores-Cruz, Z., Huang, Q., Clifford, J.M., Presting, G., et al. (2006). Identification of open reading frames unique to a select agent: *Ralstonia solanacearum* race 3 biovar 2. Mol. Plant Microbe Interact. 19, 69–79. 10.1094/MPMI-19-0069.

24. Kohlmeier, M.G., White, C.E., Fowler, J.E., Finan, T.M., and Oresnik, I.J. (2019). Galactitol catabolism in *Sinorhizobium meliloti* is dependent on a chromosomally encoded sorbitol dehydrogenase and a pSymB-encoded operon necessary for tagatose catabolism. Mol. Genet. Genomics 294, 739–755. 10.1007/s00438-019-01545-z.

25. Kohlmeier, M.G., and Oresnik, I.J. (2023). The transport of mannitol in *Sinorhizobium meliloti* is carried out by a broad-substrate polyol transporter SmoEFGK and is affected by the ability to transport and metabolize fructose. Microbiology 169, 001371. 10.1099/mic.0.001371.

26. Yamaoka, H., Yamashita, Y., Ferri, S., and Sode, K. (2008). Site directed mutagenesis studies of FAD-dependent glucose dehydrogenase catalytic subunit of *Burkholderia cepacia*. Biotechnol. Lett. 30, 1967–1972. 10.1007/s10529-008-9777-3.

27. Kerrigan, J.A., Yoshida, H., Okuda-Shimazaki, J., Temple, B., Kojima, K., and Sode, K. (2024). Improvement of substrate specificity of the direct electron transfer type FAD-dependent glucose dehydrogenase catalytic subunit. J. Biotechnol. 395, 170–179. 10.1016/j.jbiotec.2024.09.013.

28. Yamaoka, H., Ferri, S., Fujikawa, M., and Sode, K. (2004). Essential role of the small subunit of thermostable glucose dehydrogenase from *Burkholderia cepacia*. Biotechnol. Lett. 26, 1757–1761. 10.1007/s10529-004-4582-0.

29. Lowe-Power, T.M., Hendrich, C.G., von Roepenack-Lahaye, E., Li, B., Wu, D., Mitra, R., Dalsing, B.L., Ricca, P., Naidoo, J., Cook, D., et al. (2018). Metabolomics of tomato xylem sap during bacterial wilt reveals *Ralstonia solanacearum* produces abundant putrescine, a metabolite that accelerates wilt disease. Environ. Microbiol. 20, 1330–1349. 10.1111/1462-2920.14020.

30. Baroukh, C., Cottret, L., Pires, E., Peyraud, R., Guidot, A., and Genin, S. (2023). Insights into the metabolic specificities of pathogenic strains from the *Ralstonia solanacearum* species complex. mSystems 8, e00083–23. 10.1128/msystems.00083-23.

31. Geng, R., Cheng, L., Cao, C., Liu, Z., Liu, D., Xiao, Z., Wu, X., Huang, Z., Feng, Q., Luo, C., et al. (2022). Comprehensive analysis reveals the genetic and pathogenic diversity of *Ralstonia solanacearum* species complex and benefits its taxonomic classification. Front. Microbiol. 13, 854792. 10.3389/fmicb.2022.854792.

32. Good, B.H., Bhatt, A.S., and McDonald, M.J. (2025). Unraveling the tempo and mode of horizontal gene transfer in bacteria. Trends Microbiol. 33, 853–865. 10.1016/j.tim.2025.03.009.

33. Hottes, A.K., Freddolino, L., Khare, A., Donnell, Z.N., Liu, J.C., and Tavazoie, S. (2013). Bacterial adaptation through loss of function. PLoS Genet. 9, e1003617. 10.1371/journal.pgen.1003617.

34. Wen, T., Xie, P., Liu, H., Liu, T., Zhao, M., Yang, S., Niu, G., Hale, L., Singh, B.K., Kowalchuk, G.A., et al. (2023). Tapping the rhizosphere metabolites for the prebiotic control of soil-borne bacterial wilt disease. Nat. Commun. 14, 4497. 10.1038/s41467-023-40184-2.

35. Coupat-Goutaland, B., Bernillon, D., Guidot, A., Prior, P., Nesme, X., and Bertolla, F. (2011). *Ralstonia solanacearum* virulence increased following large interstrain gene transfers by natural transformation. Mol. Plant Microbe Interact. 24, 497–505. 10.1094/MPMI-09-10-0197.

36. Liu, H., Zhang, S., Schell, M.A., and Denny, T.P. (2005). Pyramiding unmarked deletions in *Ralstonia solanacearum* shows that secreted proteins in addition to plant cell-wall-degrading enzymes contribute to virulence. Mol. Plant Microbe Interact. 18, 1296–1305. 10.1094/MPMI-18-1296.

37. Fernández, E., Trillas, M.I., and Segarra, G. (2017). Increased rhizosphere populations of *Trichoderma asperellum* strain T34 caused by secretion pattern of root exudates in tomato plants inoculated with *Botrytis cinerea*. Plant Pathol. 66, 1110–1116. 10.1111/ppa.12668.

38. Aramaki, T., Blanc-Mathieu, R., Endo, H., Ohkubo, K., Kanehisa, M., Goto, S., and Ogata, H. (2020). KofamKOALA: KEGG ortholog assignment based on profile HMM and adaptive score threshold. Bioinformatics 36, 2251–2252. 10.1093/bioinformatics/btz859.

39. Chen, T., and Guestrin, C. (2016). XGBoost: A Scalable Tree Boosting System. In Proceedings of the 22nd ACM SIGKDD International Conference on Knowledge Discovery and Data Mining KDD ‘16. (Association for Computing Machinery), pp. 785–794. 10.1145/2939672.2939785.

40. Lees, J.A., Galardini, M., Bentley, S.D., Weiser, J.N., and Corander, J. (2018). pyseer: A comprehensive tool for microbial pangenome-wide association studies. Bioinformatics 34, 4310–4312. 10.1093/bioinformatics/bty539.

41. Yan, J., Lin, N., Wang, X., Chen, X., Wang, H., Lin, Q., Zhou, X., Zhang, L., and Liao, L. (2022). Markerless gene deletion in *Ralstonia solanacearum* based on its natural transformation competence. Front. Microbiol. 13, 977580. 10.3389/fmicb.2022.977580.

42. Emms, D.M., Liu, Y., Belcher, L., Holmes, J., and Kelly, S. (2026). OrthoFinder: Improved phylogenetic orthology inference with enhanced accuracy and scalability. Nat. Methods 23, 1327–1333. 10.1038/s41592-026-03126-6.

43. Katoh, K., and Standley, D.M. (2013). MAFFT multiple sequence alignment software version 7: Improvements in performance and usability. Mol. Biol. Evol. 30, 772–780. 10.1093/molbev/mst010.

44. Capella-Gutiérrez, S., Silla-Martínez, J.M., and Gabaldón, T. (2009). trimAl: A tool for automated alignment trimming in large-scale phylogenetic analyses. Bioinformatics 25, 1972–1973. 10.1093/bioinformatics/btp348.

45. Borowiec, M.L. (2016). AMAS: A fast tool for alignment manipulation and computing of summary statistics. PeerJ 4, e1660. 10.7717/peerj.1660.

46. Nguyen, L.-T., Schmidt, H.A., von Haeseler, A., and Minh, B.Q. (2015). IQ-TREE: A fast and effective stochastic algorithm for estimating maximum-likelihood phylogenies. Mol. Biol. Evol. 32, 268–274. 10.1093/molbev/msu300.

47. Letunic, I., and Bork, P. (2024). Interactive tree of life (iTOL) v6: Recent updates to the phylogenetic tree display and annotation tool. Nucleic Acids Res. 52, W78–W82. 10.1093/nar/gkae268.

48. Gilchrist, C.L.M., Booth, T.J., van Wersch, B., van Grieken, L., Medema, M.H., and Chooi, Y.-H. (2021). cblaster: A remote search tool for rapid identification and visualization of homologous gene clusters. Bioinform. Adv. 1, vbab016. 10.1093/bioadv/vbab016.

49. Simão, F.A., Waterhouse, R.M., Ioannidis, P., Kriventseva, E.V., and Zdobnov, E.M. (2015). BUSCO: Assessing genome assembly and annotation completeness with single-copy orthologs. Bioinformatics 31, 3210–3212. 10.1093/bioinformatics/btv351.

50. Waterhouse, A., Bertoni, M., Bienert, S., Studer, G., Tauriello, G., Gumienny, R., Heer, F.T., de Beer, T.A.P., Rempfer, C., Bordoli, L., et al. (2018). SWISS-MODEL: Homology modelling of protein structures and complexes. Nucleic Acids Res. 46, W296–W303. 10.1093/nar/gky427.

51. Gilchrist, C.L.M., and Chooi, Y.-H. (2021). clinker & clustermap.js: Automatic generation of gene cluster comparison figures. Bioinformatics 37, 2473–2475. 10.1093/bioinformatics/btab007.

52. Ives, A.R., and Garland, T. (2010). Phylogenetic logistic regression for binary dependent variables. Syst. Biol. 59, 9–26. 10.1093/sysbio/syp074.

53. Tung Ho, L.S., and Ané, C. (2014). A linear-time algorithm for Gaussian and non-Gaussian trait evolution models. Syst. Biol. 63, 397–408. 10.1093/sysbio/syu005.

54. Bürkner, P.-C. (2017). brms: An R package for Bayesian multilevel models using Stan. J. Stat. Softw. 80, 1–28. 10.18637/jss.v080.i01.

55. Revell, L.J. (2012). phytools: An R package for phylogenetic comparative biology (and other things). Methods Ecol. Evol. 3, 217–223. 10.1111/j.2041-210X.2011.00169.x.

56. Sharma, P., Johnson, M.A., Mazloom, R., Allen, C., Heath, L.S., Lowe-Power, T.M., and Vinatzer, B.A. (2022). Meta-analysis of the *Ralstonia solanacearum* species complex (RSSC) based on comparative evolutionary genomics and reverse ecology. Microb. Genomics 8, 000791. 10.1099/mgen.0.000791.

